# ERECTA signaling controls the timing of Arabidopsis Guard Cell maturation at the embryonic leaf tip

**DOI:** 10.64898/2026.08.10.743849

**Authors:** Yadhusankar Sasidharan, Vijayalakshmi Suryavanshi, Pablo González-Suárez, Steffi Zimmermann, Sandra Richter, Friederike Hauschild, Agneta Larsina Timpe, Sarah-Karina Loosen, Margot E. Smit

## Abstract

While cell identities are established early during embryogenesis, these cells remain immature until germination, and the mechanisms enforcing this developmental pause are poorly understood. Embryonic stomatal cells provide a model to study this pause as the stomatal transcription factor FAMA, normally sufficient for Guard Cell maturation in seedlings, can not drive maturation in the Arabidopsis embryo. Here we show that FAMA’s ability to drive maturation depends on leaf polarity and adaxial stomatal cells can progress further in their lineage. We next find that ERECTA-family receptor signaling, which controls stomatal patterning, also suppresses embryonic stomatal maturation. In *er erl1 erl2* mutants, cell pairs at the cotyledon tip acquire characteristics of maturing guard cells: cell wall reinforcement, pore-associated thickening, and expression of late lineage markers as identified by whole embryo transcriptomics. This precocious maturation however remains incomplete: many GC markers remain absent, and cells lack an open pore and mature vacuoles. Genetic analysis shows that partial maturation requires but is not limited by low levels of FAMA. Restriction of maturation to the cotyledon tip correlates with locally elevated ERECTA-family receptor abundance, while high auxin appears dispensable for this. Finally, we show that EPFL-ER signaling mediates leaf tip Guard Cell size postembryonically as well. Altogether, we identify ERECTA signaling as a local brake on embryonic stomatal cell maturation, discovering another way to push precocious stomatal cell maturation that results in a complex, partially mature cell state that provide insights into the limitations on cell embryonic cell maturation.

## Introduction

Plant embryogenesis relies on tightly orchestrated cell division and specification leading to the formation of embryonic organs containing pre-patterned cells ready to continue development upon germination. Mature, functional cell types are absent during embryogenesis and the mechanisms that control cell maturation timing remain unknown^1,2^.

In Arabidopsis, the majority of embryonic cell identities have been laid down by heart stage^3,4^. Leaf-specific epidermal identities are specified afterwards with stomatal precursors first forming at bent cotyledon stage^5^. Following specification, embryonic cells divide and progress to intermediate states and express mid-lineage reporters but arrest before maturation^6,7^. This pause of cell development during seed maturation coincides with programs that coordinate nutrient storage and dormancy^4,8^. The mature embryo in the dry seed thus contains specified but immature cells that become functional rapidly upon germination^9,10^.

In the seedling, stomata are produced through a series of coordinated divisions and cell identity changes that are coordinated by three main basic Helix-Loop-Helix (bHLH) Transcription Factors (TFs)^11–13^. First, SPEECHLESS (SPCH) promotes Asymmetric Cell Division (ACD) of protodermal cells, with the smaller daughter cell becoming a stomatal Meristemoid^14^. After several rounds of ACDs^15^, MUTE induces the transition to Guard Mother Cell (GMC) state and changes cell divisions to produce symmetrical Guard Cell (GC)l pairs^16^. Finally, FAMA inhibits further divisions and promotes GC maturation, resulting in a complex of two cells surrounding a pore^17^. SPCH, MUTE, and FAMA each require the presence of heterodimerization partners SCRM/ICE1 (SCREAM/INDUCER OF CBF EXPRESSION1) or SCRM2 for their function^18^.

During the final step of stomatal development, FAMA is necessary and sufficient for stomatal maturation. Accordingly, *fama* mutants are devoid of functional GCs and ectopic FAMA induces GC maturation and pore formation in aerial epidermal cells as well as shape and gene expression changes in other cell types^17^. Functional stomatal complexes have distinct physical features including reinforced cell walls that are strengthened around their central pore that can open, an external cuticular ledge, symplastic isolation via plasmodesmatal closure, and large, dynamic vacuoles enabling pore closure^19–24^. Maturing and mature GCs also express stage-specific genes. During maturation, FAMA induces expression of WASABIMAKER (WSB), followed by STOMATAL CARPENTER1 (SCAP1), each of which play key roles in GC functionalization^25^. Later on, GCs express TFs such as MYB60^26^ as well as ion channels including SLAC1 (SLOW ANION CHANNEL-ASSOCIATED1), GORK (GUARD CELL OUTWARDLY RECTIFYING K⁺ CHANNEL) and KAT1/2^27–30^. Recently, integrative analysis of published single cell RNA-seq datasets identified transcripts specific to maturing and mature GCs^31^.

Stomatal patterning is carefully regulated by signaling within the epidermis and between cell layers^11^. Perception of EPIDERMAL PATTERNING FACTOR (EPF) and EPF-LIKE (EPFL) mobile peptides takes place by ERECTA (ER) receptors with or without the TOO MANY MOUTHS (TMM) co-receptor, SOMATIC EMBRYOGENESIS RECEPTOR KINASE (SERK)^32^ coreceptors, and with the membrane-attached cytoplasmic kinases BRASSINOSTEROID-SIGNALING KINASES 1 (BSK1) and BSK2^33^. Perception triggers a Mitogen-Activated Protein Kinase (MAPK) phosphorylation cascade involving YODA(YDA)-MKK5/6-MPK3/6 which in stomatal cells results in the phosphorylation and degradation of SPCH and its heterodimerization partners^34–41^. Thus, EPF-ER signaling inhibits stomatal initiation and controls stomatal numbers and spacing with exact outcomes depending on local differences in peptide and receptor production^42,43^. While MAPK signaling was shown to directly destabilize SPCH and SCRM1/2, direct effects remain unknown for MUTE and FAMA^37,44^. However, EPF1 perception by ERECTA-LIKE1 (ERL1) reduces MUTE expression and protein levels^45^ and MAPK signaling can have a positive effect on the final stages of stomatal development^44^. Thus, EPF-ER signaling affects all stages of stomatal development and indeed, higher order ER mutants produce more mature stomata. Absence of ER signaling might even allow bypassing key regulators such as MUTE, with *er erl1 erl2 mute* seedlings producing occasional stomata^46^, suggesting that other stomatal bHLHs could drive additional identity transitions^46–49^. Finally, *er erl1 erl2* mutants already show additional mature GC complexes two days after germination, suggesting that ER signaling controls maturation speed upon germination^50^ and prompting us to investigate a potential role in embryonic maturation.

Many studies on stomatal development have used cotyledons as their model tissue. Cotyledons differ from other leaf types in their developmental origin and function as they play key roles in energy storage and desiccation tolerance during embryogenesis^4,8^. In the embryonic stages of cotyledon development, stomatal cells do not become functional^9^. Recent work suggests that stomatal maturation is being prevented since misexpression of FAMA cannot promote it^5^. Instead, embryonic FAMA is limited to inducing cell size increase and expression of FAMA’s direct target *WSB*, indicating that FAMA’s ability to induce cell state progression is regulated during this developmental stage.

Here, we ask what factors limit FAMA’s ability to induce embryonic GC maturation. We find leaf polarity limits FAMA ability while embryonic levels of its heterodimerization partner SCRM do not. We find that ER signaling prevents embryonic stomatal maturation specifically at the cotyledon tip. These precociously differentiated cells in *er erl1 erl2* undergo partial maturation prior to germination, as evidenced by transcriptomic and morphological features. Finally, we investigate the mechanisms that might be responsible for spatially limiting this maturation to the cotyledon tip.

## Results

### Leaf polarity impacts FAMA’s ability to promote GC state

To further characterize the cell state induced by embryonic FAMA, we used modified Pseudo-Schiff Propidium Iodide (mPS-PI) staining followed by the construction of 2.5D meshes of epidermal surfaces^51,52^. In mature EPF2p::FAMA embryos, single stomatal cells on the abaxial surface increase in size but lack further morphological changes^5^. In contrast, pairs of stomatal cells on the adaxial surface gain intense PI staining indicating cell wall reinforcement^51^ (**Figure1A-D’**). In addition, the GC marker SCAP1^53^ is expressed only in pairs of adaxial stomatal cells, indicating that embryonic GC progression depends on leaf polarity (**Figure1E**,**S1A**).

**Figure 1:**
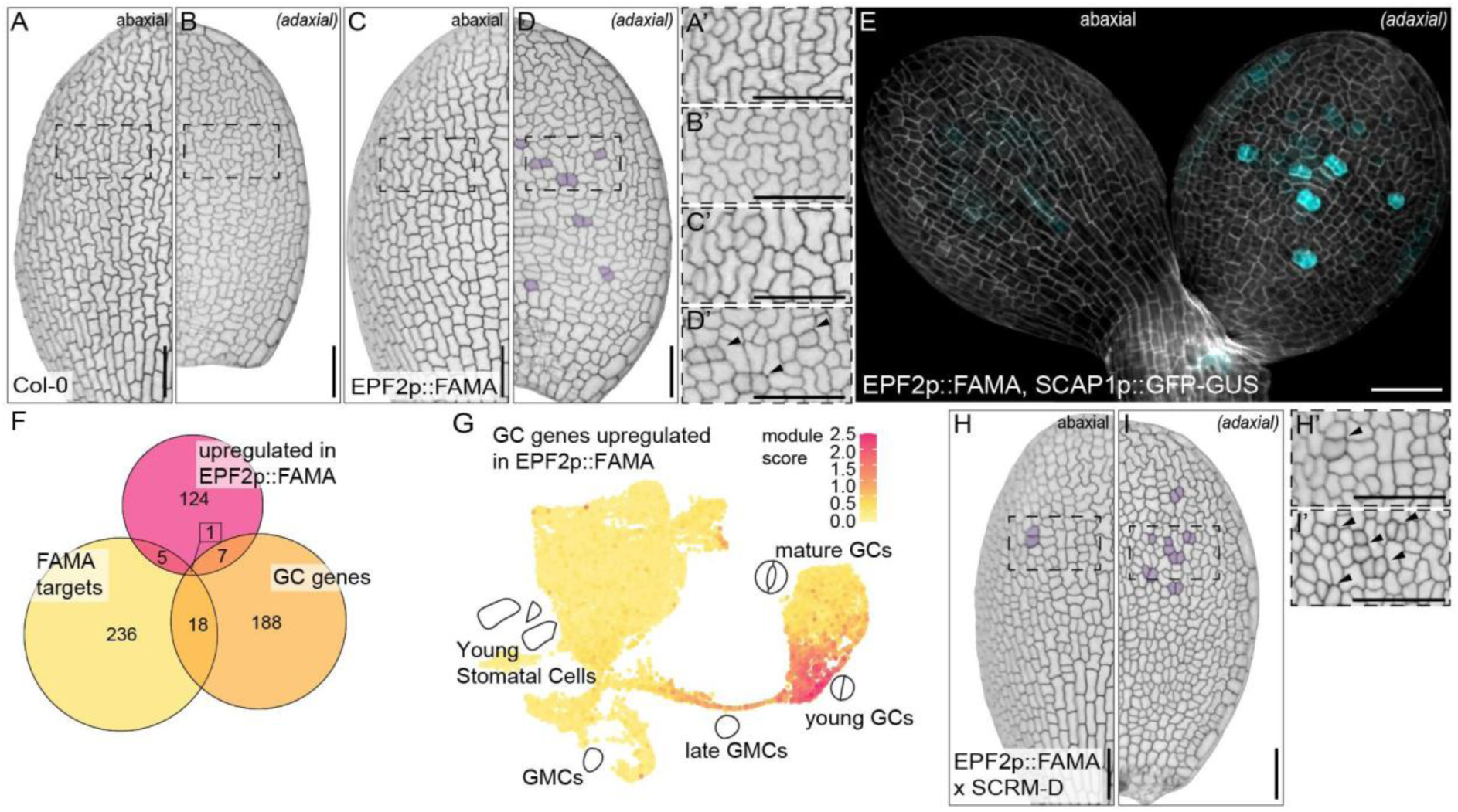
Progression of stomatal cell maturation as a result of FAMA activity. (A-D) Cotyledon surface images of mature embryos. Abaxial (A,C) and adaxial (B,D) surfaces of Col-0 (A-B) and *EPF2p::FAMA* (C-D) mature embryos from dry seeds. (A’-D’) Zoomed ins showing with arrowheads highlighting abnormal cell morphology. (E) Expression of a SCAP1p::GFP-GUS in *EPF2p::FAMA* embryos. (F) Venn diagram showing overlap between genes upregulated in *EPF2p::FAMA* embryos with FAMA targets and GC genes. (G) GC genes up-regulated in EPF2p::FAMA embryos plotted on a stomatal development UMAP plot^55^ highlighting their enrichment in young GCs. (H-I) Cotyledon surface images of SCRM-D, *EPF2p::FAMA* mature embryos. (H’-I’) Zoomed in images of surface highlights showing abnormal cell morphology highlighted with arrowheads. Images in A-E and H-I were constructed with 2.5-D processing of image stacks of PI (A-D, H-I) or R2200 and GFP signals (E) in MorphographX. Scale bars indicate 50 μm. Purple highlights in A-D,H-I indicate pairs of cells with increased PI iodide staining.

To characterize the degree of maturity in these cells, we generated transcriptomes of Bent Cotyledon stage embryos and identified 137 genes upregulated in EPF2p::FAMA embryos compared to Col-0 (**Figure1F**). Among these, Gene Ontology (GO) term analysis indicated enrichment in response to hypoxia and metabolic reprogramming (**FigureS2A**,**TableS1-2**). Since stomatal GO terms are sparsely populated, we decided to test for enrichment in GC maturation using more complete sets of GC-specific markers and FAMA targets. A set of 586 mature stomatal genes had previously been curated using snRNA-seq datasets^31^. We narrowed this set down to 214 GC-specific genes by stringently filtering against early stomatal expression and expression in other lineages (**TableS3**). In addition, we compiled a list of 260 FAMA target genes from previous datasets^25,54^(**TableS4**). Using these we found that 5 out of 137 upregulated genes are targets of FAMA and 7 are GC-specific (Excluding FAMA, **Figure1F**,**TableS5**), representing statistically significant overrepresentation in both categories (**FigureS2B**)(hypergeometric test, p=0.0003 and p=0.02, respectively). A closer look at the upregulated GC genes revealed that they are mainly expressed in young GCs during post-embryonic stomatal development (**Figure1G**)^55,56^, suggesting that embryonic FAMA can promote stomatal progression but acquisition of only young GC features. These are normally absent from the embryo, indicating real but limited progression.

### SCRM presence and activity do not limit FAMA-driven GC maturation

FAMA requires a heterodimerization partner for its activity. While both SCRM and SCRM2 are expressed during embryogenesis^5^, their stability and activity might be altered. To determine whether dimerization partner activity limits embryonic maturation potential, we observed F1 EPF2p::FAMA *scrm-D* embryos and found that they produce more stomatal cells in general and slightly more specific GCs specifically as seen by SCAP1+ cells (**FigureS1B-D**). However, GCs do not appear to progress further and lack pore formation (**Figure1H-I**), indicating that additional factors still limit FAMA’s ability to promote GC maturation.

### ERECTA triple mutants produce embryonic GCs with maturation features

ER signaling regulates stomatal numbers, spacing and maturation^34,41,57^, and *er erl1 erl2* seedlings show fast GC maturation upon germination^50^. To determine when stomata mature upon germination in the absence of ER signaling, young seedlings of Col-0 and segregating *er erl1 erl2* (see Star Methods on details of two lines) were collected every 12 hours. In Col-0, mature stomata are first found across the cotyledon surface 48 hours after the transfer to light, at which point *er erl1 erl2* seedlings contain many clustered mature stomata (**Figure2A-D,S3,S4**). Looking earlier, *er erl1 erl2* seedlings already have mature GCs at 12h at the cotyledon tip with non-tip GCs quickly following at 24-36h (**Figure2A-D,S3,S4**). This initial maturation is present before the cotyledon emerges from the seed coat, but in addition *erl1 erl2* seedlings have faster overall development: they germinate earlier and have faster root growth in the initial days (**Figure2E**).

**Figure 2:**
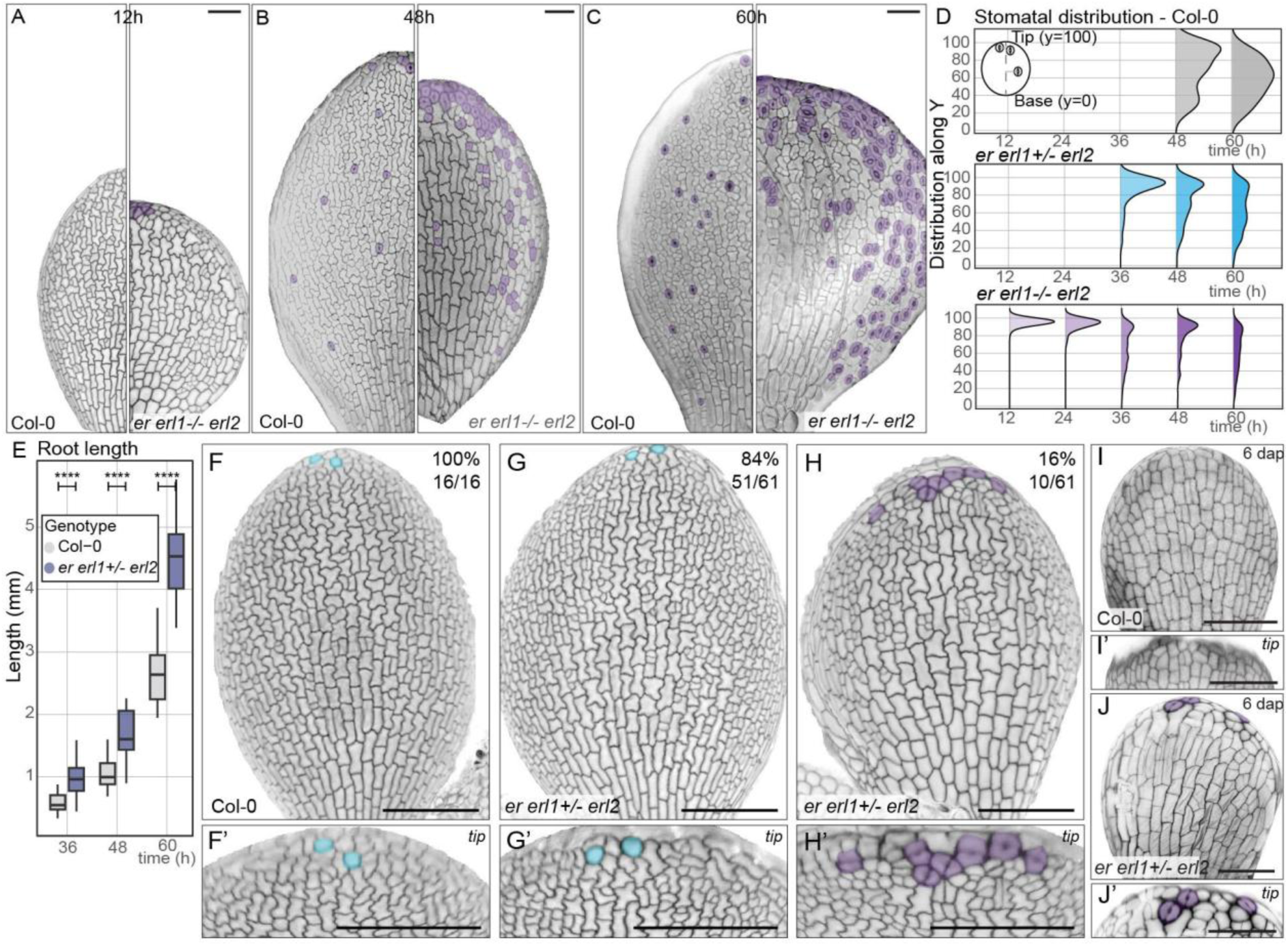
Mutants lacking ER signaling produce precocious stomata starting during embryogenesis. (A-D) Epidermal surface images of Col-0 (left) and *er erl1-/- erl2* (right) germinating seedlings at 12 (A), 36 (B), and 60 (C) hours after being placed into light. (D) Quantification of stomatal maturation timing and location upon germination. n=2-6 leaves. (E) Quantification of root length of Col-0 and segregating *er erl1+/- erl2* upon germination. **** Indicate p-values below 0.0001, n=25 seedlings. (F-H’) Embryonic epidermis images of Col-0 (F-F’) and *er erl1+/- erl2* (G-H’), lineage precursors at the tip highlighted in blue and tilted zoom-ins in. (I-J’) 6DAP abaxial epidermis images for Col-0 (I-I’) and *er erl1+/- erl2* (J-J’). Images in A-C and F-J’ were constructed with 2.5-D processing of image stacks of PI signal in MorphographX. Across all images, purple highlights indicate GCs and GC-like cells. Scale bars in F-H indicate 100 µm, other scale bars indicate 50 µm.

As GCs are present before cotyledon emergence, we asked whether GCs mature during embryogenesis in *er erl1 erl2*. 16% and 28% of embryos from segregating *er erl1/+ erl2* and *er/+ erl1 erl2* seeds showed pairs of cells with GC-like morphology at the cotyledon tip (**Figure2F-H,S5**). GC-like cell pairs had increased PI staining indicating cell wall reinforcement as well as central cell wall thickening at the future pore site (**Figure2H-H’**). Remarkably, we found these features as early as 6 days after fertilization, at Torpedo stage, indicating rapid progression to cell maturation upon stomatal specification compared to Col-0 (**Figure2I-J’**). Thus, ER signaling affects tip-specific GC maturation during embryogenesis.

Based on cell shape and cell wall thickening, we hypothesized that GCs of *er erl1 erl2* embryos are underway to cell maturation. To further investigate their cell state, we introduced reporters for several late stomatal lineage markers. In wild-type, expression of *SDD1*, *SLL3*, and *FAMA* starts in late GMCs post-embryonically^17,48,58^ and is absent during embryogenesis. In contrast, we detected expression of all three markers in cells at the tip of *er erl1 erl2* embryos, indicating that stomatal cells are progressing to later stages (**Figure3A-C**). The next marker, *SCAP1* is normally expressed in maturing GCs^25,53,54^, but is not expressed in *er erl1 erl2* embryos, indicating that these cells lack some crucial aspects of the GC maturation program.

**Figure 3:**
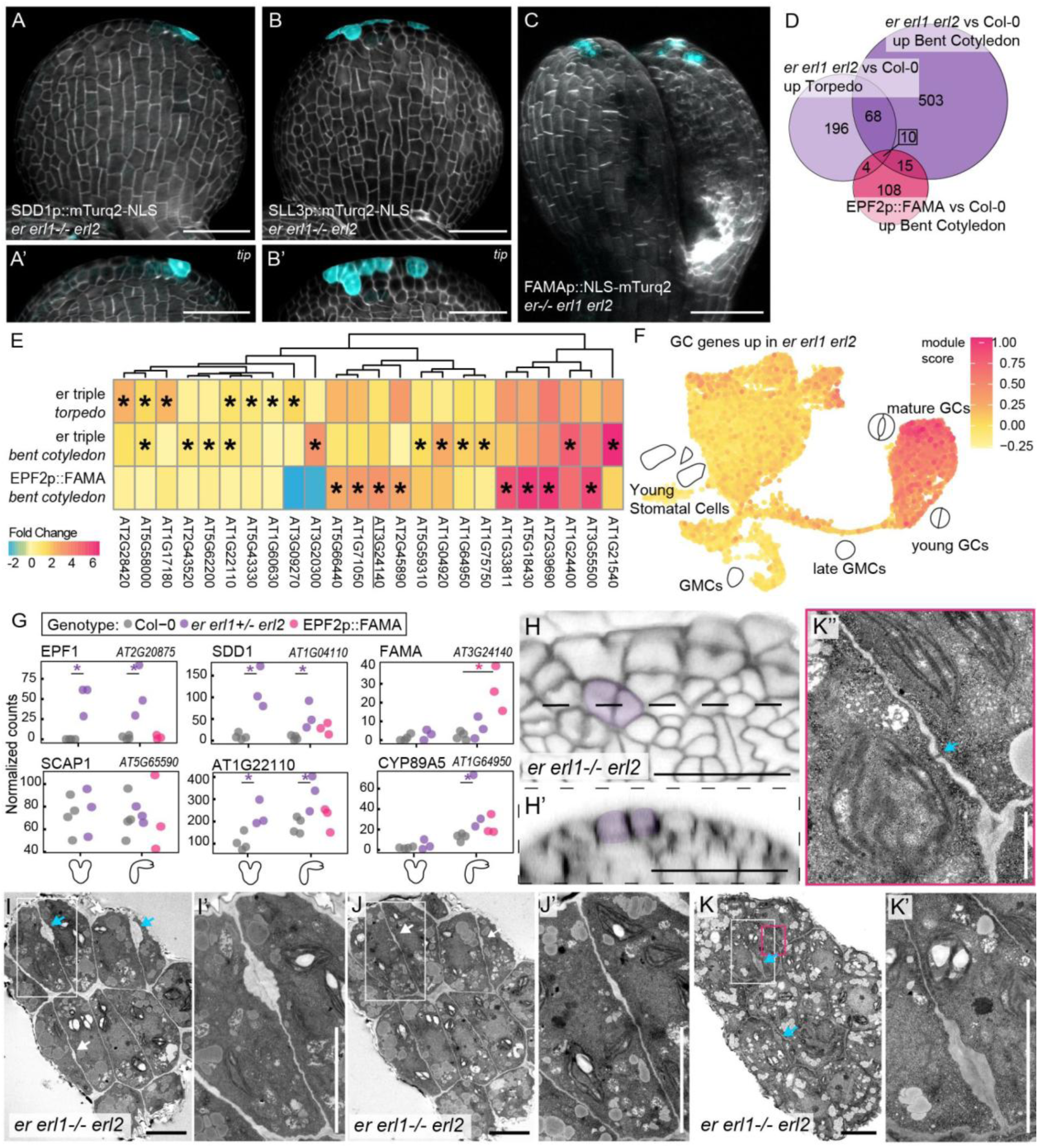
GCs in *er erl1 erl2* embryos are lack key maturation features. (A-C’) Expression SDD1, SLL3, and FAMA is specific to the cotyledon tip. (D) Venn diagram showing genes upregulated in EPF2::FAMA or *er erl1 erl2* embryos. (E) Heat map of differentially expressed GC genes in *er erl1 erl2,* and EPF2::FAMA with respect to WT. (F) GC genes up-regulated in *er erl1 erl2* embryos plotted on a stomatal UMAP plot^55^ highlighting their enrichment in mature GCs. (G) Expression of specific GC genes across transcriptomic samples. (H) Close-up of GC-like cells at the tip of *er erl1 erl2* embryos with (H’) showing a transverse section indicating increased PI signal at the pore formation site. (I-K’’) TEM images showing pore cell wall thickening at the site of pore formation, indicated by blue arrows. (J-J’) Subsequent sections confirm the absence of the pore (white arrows) but show additional thickening at the cell base. (K-K’) Deeper sectioning reveals reappearing cell wall thickening at putative pore site. (K’’) Image showing presence of plasmodesmos between a GC and its neighbour. Images in A-C and H were constructed with 2.5-D processing of image stacks of R2200 and fluorescence signals (A-C) or PI (H) in MorphographX. Scale bars in A-C&H indicate 100 µm, in I-K’ 5 µm, and in K’’ 1 µm.

To characterize GC maturation in *er erl1 erl2* embryos, we generated transcriptomes of *er erl1 erl2* embryos at Bent Cotyledon and Torpedo stages. 281 and 596 genes were upregulated in *er erl1 erl2* at Torpedo and Bent Cotyledon stages, respectively (**Figure3D**,**TableS6,S7**). Among these, we found early to mid-stomatal genes such as EPF2, TMM, and MUTE, which are normally already expressed in the embryo aligning with *er erl1 erl2* embryos producing more stomatal precursors. GO term analysis identified “shoot system morphogenesis” and “developmental growth in morphogenesis” as enriched processes among upregulated genes (**FigureS6C-D**,**TableS8**), reflecting altered tissue morphology in *er erl1 erl2* mutants^57^. To specifically test for the presence of GC maturation, we utilised our previously curated gene sets. Despite a multitude of cell types being altered in *er erl1 erl2* mutants^40,50,59^, GC genes are overrepresented among upregulated genes at Torpedo and Bent Cotyledon stages (7 and 11 of 214 GC genes upregulated, respectively)(**Figure3E,S5E-H**,**TableS9**)(hypergeometric test, p=0.02 and p=0.03, respectively). However, several well-known GC genes, including *WSB*, *SCAP1*, *GORK*, and *SLAC1* are not significantly upregulated (**Figure3G,S5F-G**), indicating the absence of key aspects of GC identity. Closer examination of GC-related genes upregulated in *er erl1 erl2* using available scRNA-seq data of the stomatal lineage^55^ revealed they are predominantly expressed in late stages of GC maturation post-embryonically^55,56^. Together, we conclude that cells in *er erl1 erl2* embryos gain features maturing stomata but that key features of mature GCs remain absent.

While gene expression is one indicator of cell state, GCs’ distinct physical features are additional indicators of maturity and functionality. Propidium iodide staining indicated cell wall thickening indicative of pore formation (**Figure3H**). Using Transmission Electron Microscopy (TEM) we find that maturing GCs in *er erl1-/- erl2* embryos lacked large vacuoles and an open pore, and retain plasmodesmata, indicating incomplete stomatal maturation and lack of functionality (**Figure3I-K’,S7**). However, the characteristic cell wall thickening at the future pore site is present between pairs of cells. Interestingly, this thickening appears most extreme near the upper and lower cell surfaces but relatively minimal in between. Thus, *er erl1-/- erl2* embryonic GCs possess developing pores while lacking large vacuoles and cytoplasmic isolation.

### Canonical ER signaling controls embryonic GC maturation together with FAMA

The ER signaling cascade is well-characterized and mutants of actors downstream of ER show similar stomatal patterning phenotypes^11^. We next sought to explore underlying mechanisms and confirm the role of the canonical ER signaling pathway in preventing precocious maturation. However, since embryos any remaining ER homolog showed no embryonic GCs, we believed that these can only form in the complete absence of ER signaling, something that often results in sterility or lethality when the required homologs of downstream components are absent^34,60^.

Upstream of ER, EPF1 and EPF2 peptides regulate stomatal patterning (**Figure4A**)^11,36,61^. Similarly, during embryogenesis, *epf1 epf2* mutants produce additional stomatal cells (**FigureS8A-A’**). However, maturing GCs remain absent, indicating that prevention of embryonic GC maturation depends on perception of different or additional EPF/EPFL peptides. Similarly, when considering *bsk1 bsk2* mutants^33^, we find an increase in stomatal cells but only 1 out of 31 *bsk1 bsk2* individuals showed signs of GC formation (**Figure4B,S8B-B’**). Mutant embryos of the final component investigated, the key MAPKKK YDA^35^, show smaller and rounder cotyledons with increased numbers of stomatal precursors and several embryos (5 out of 30) also have maturing GCs, though there are fewer complexes per embryo and pore cell wall thickening was less prominent (**Figure4C-D’,S4C**). Overall, our findings indicate that any remaining ER signaling activity will prevent embryonic GC formation but that this process is indeed inhibited by canonical ER.

In our *er erl1 erl2* transcriptomes, we found that *FAMA* expression is slightly increased compared to wild-type, but that this trend was barely detectable and not significant (**Figure3G**). A transcriptional *FAMA* reporter indicates that *FAMA* is specifically and clearly expressed in the maturing GCs of *er erl1 erl2* (**Figure3C**). To determine whether FAMA is required for GC maturation, we generated *er/+ erl1/+ erl2 fama/+* quadruple mutants and imaged individuals with smaller, rounded cotyledons characteristic of *er erl1 erl2*. We found that a large number of these embryos (6 out of 10) no longer formed pairs of GCs (**Figure4E-F**), indicating that those low levels of FAMA are required for the GC maturation processes seen in *er erl1 erl2*.

**Figure 4:**
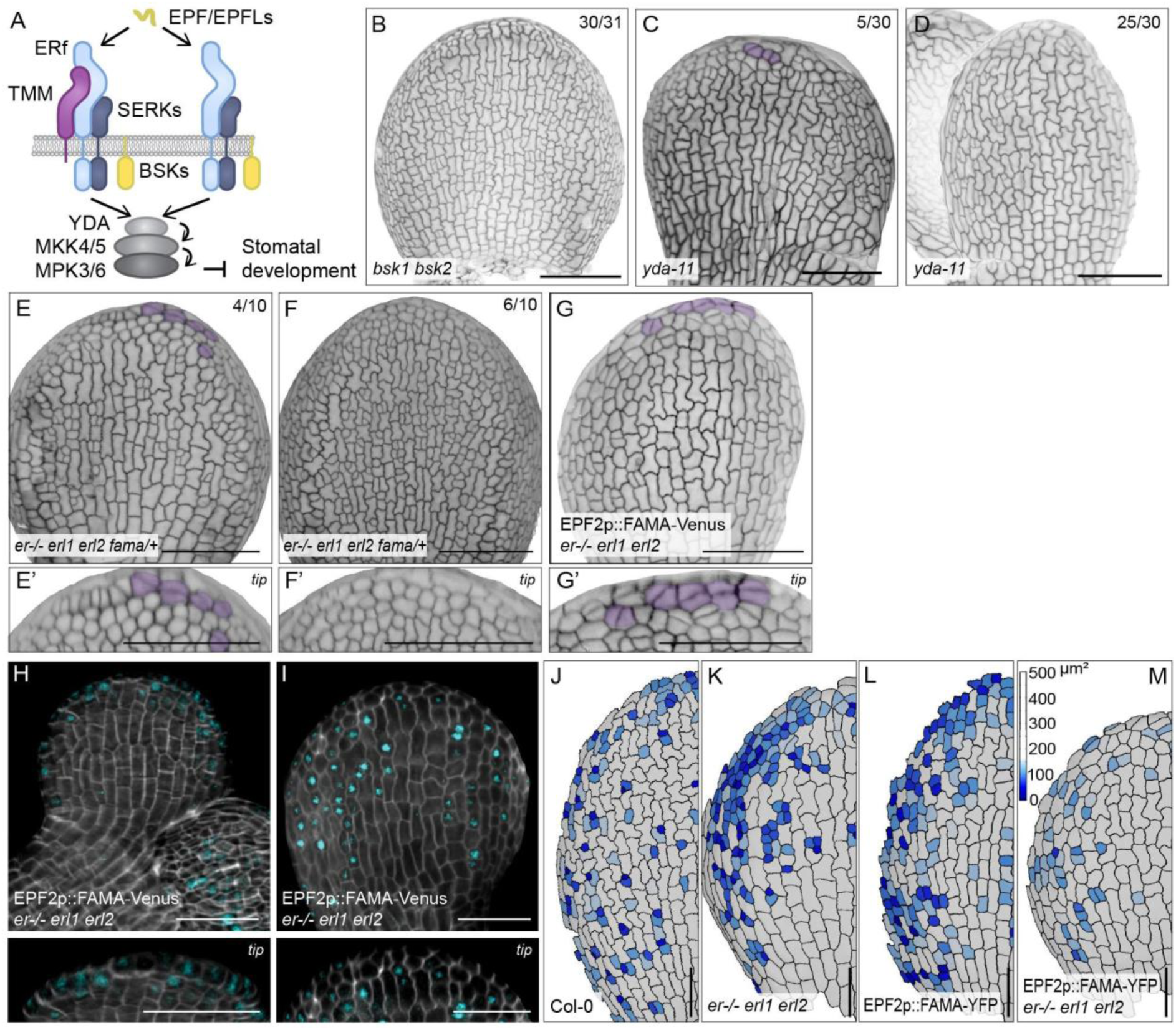
ER signaling and FAMA affect separate aspects of GC maturation. (A) Schematic of canonical ER signaling affecting stomatal development. (B) Embryonic epidermis images of *bsk1 bsk2* mutants, (C-D) *yda-11* mutants, (E-F’) *er+/- erl1+/- erl2 fama+/-* mutants and (G-G’) EPF2::FAMA *er erl1 erl2* lines. (H-I) Developing embryo showing FAMA-Venus expression in EPF2::FAMA *er erl1 erl2* epidermal cells but signal absence in maturing GCs. (J-M) Heatmaps showing cell area distribution in Col-0 (J), *er+/- erl1 erl2* (K), EPF2::FAMA (L) and EPF2::FAMA *er+/- erl1 erl2* (M). Scale bars in A-G indicate 100 µm, in H-M 50 µm.

Both FAMA presence and ERECTA absence allow for embryonic stomatal cells to gain features of maturing GCs. Interestingly, these features only show little overlap: FAMA induces early GC-enriched gene expression and stomatal cell size increase across the leaf, while *er erl1 erl2* embryos express late stomatal genes and contain maturing GCs specifically at the tip. We wondered whether low levels of FAMA limited GC maturation *er erl1 erl2* mutants and whether the presence of ER signaling prevented further GC maturation upon embryonic FAMA expression. Surprisingly, EPF2p::FAMA *er erl1 erl2* embryos show the two distinct phenotypes separately: embryos have tip-located partially mature pairs and surface-located enlarged cells (**Figure4G-M**). However, FAMA-Venus cannot be detected in all tip-located pairs (**Figure4H-I’**), indicating that either FAMA is absent or only shortly expressed under the EPF2 promoter as cells rapidly move to maturation. Thus, we find indications that the lack of full GC maturation is not merely due to FAMA absence or ER presence but might involve additional mechanisms.

### What makes GCs at the cotyledon different?

High auxin signalling is one of the factors that sets apart the leaf tip and this is unaltered in *er erl1 erl2* embryos^50^. To test whether high auxin limits the location and degree of GC maturation, we cultured developing seeds with and without the synthetic auxin 2,4-D. While auxin treatment was effective as evidenced by increased vascular development in the cotyledon (**FigureS9A-B**), it did not result in the production of additional or more mature GC complexes (**Figure5A-B,S9D**). Thus, it appears that embryonic GC maturation is not restricted by auxin levels.

**Figure 5:**
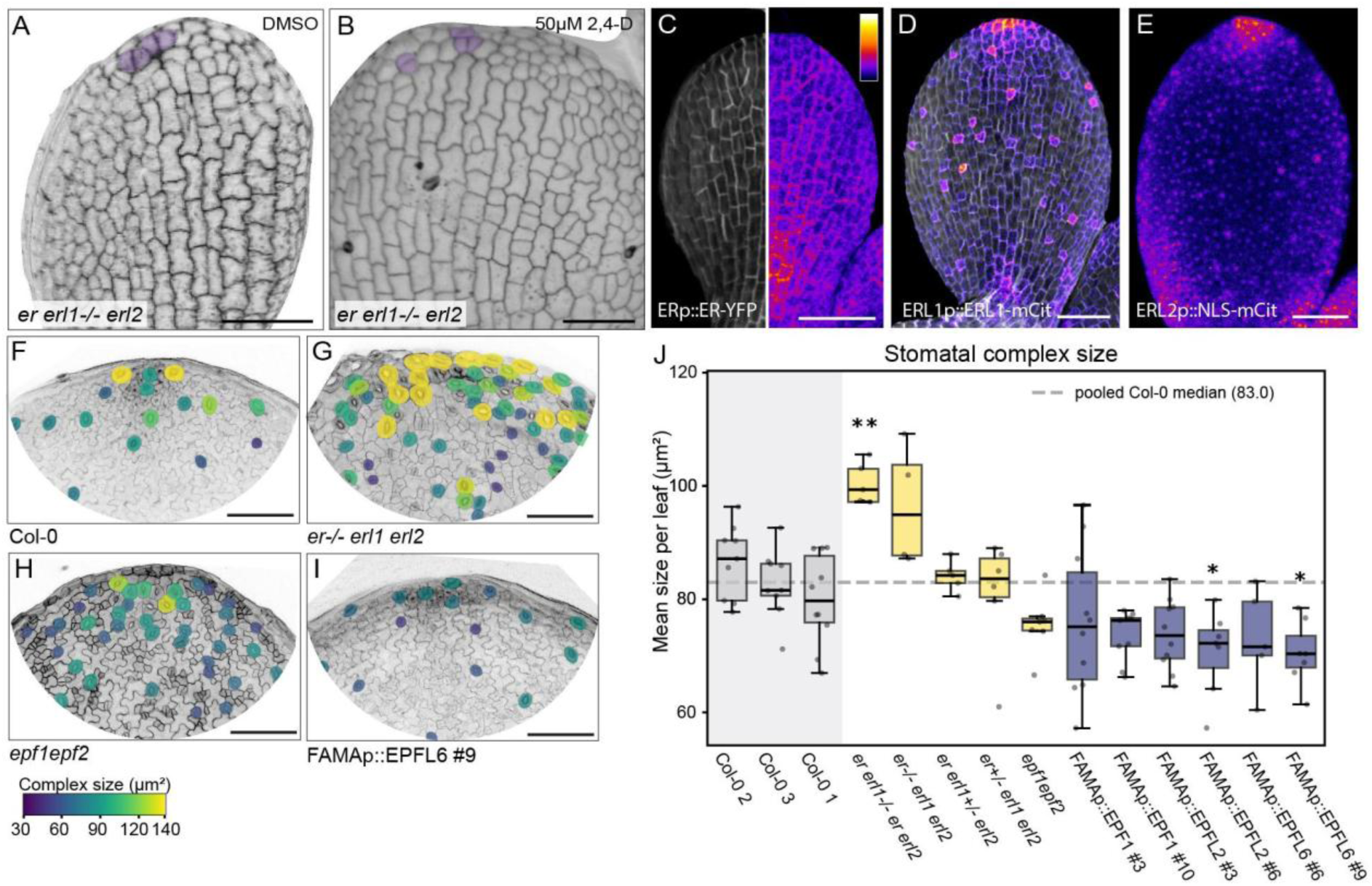
Leaf tip-specific GC features and mechanisms. (A-C) Embryonic expression of ER receptors in the cotyledon, ERp::ER-YFP (A), ERL1p::ERL1-mCit (B) and ERL2p::NLS-mCit (C). Images were processed with MorphographX, extracting surface signal from image stacks of R2200 and fluorescence signals. (D-E) Embryonic surface images of embryos treated with or without 2,4-D in seed culture experiments. (F-I) Leaf tip images of 4 DAG seedlings with stomata within 200 µm of the leaf tip manually segmented and colored by complex size for Col-0 (F), *er erl1 erl2* (G), *epf1 epf2* (H), and FAMAp::EPFL6. (J) Quantification of mean stomatal size across different genotypes (n= 4-10 leaves)(* indicates p<0.05, ** indicates p<0.01 as tested using Dunnett’s comparing test samples to the pooled WT, Col-0). Scale bars in A-E indicate 50 µm, in F-I 100 µm.

The restriction of GC maturation to the cotyledon tip is striking and led us to question what features of the leaf tip affect the GC maturation programme. ER itself is not more abundant in the tip and might even be slightly depleted in the underlying mesophyll (**Figure5C,S9C**), but abundance of its homologs is higher at the cotyledon tip (**Figure5B-C**). Reporters for ERL1 and ERL2 revealed increased expression and protein, respectively, in epidermal cells at the cotyledon tip (**Figure5B-C**), indicating that the tip-specificity might partly rely on spatial distribution of ER receptors.

The tips of leaves are characterized by their regional hydathode identity and hydathode GCs are larger than other GCs as well as unable to fully close, indicating they undergo specific maturation steps^62–64^. We need hydathode GC-specific reporters to determine whether maturing GCs in *er erl1 erl2* embryos are specifically hydathode GCs. As such reporters are not yet known, we attempted to identify such genes by combining several stomatal and hydathode transcriptomic studies^49,55,63,65,66^ but were unable to generate a true hydathode GC-specific reporter. Therefore we could not determine whether the maturing GCs in *er erl1 erl2* embryos are specifically hydathode GCs.

While we could not identify what transcriptomic features set apart hydathode GCs, we wondered whether ER signaling might specifically contribute to the unique features, mainly size, of hydathode GCs. In *er erl1 erl2* seedlings, stomata are clustered and large^40,42,45^ but it was unclear whether the size increase was the result of clustering or a direct result of ER signaling. We measured stomatal size in the leaf tip area and confirmed that *er erl1 erl2* individuals had larger stomata (**Figure5F,G,J**). However, we find that similar clustering in *epf1 epf2* did not result in a similar size increase (**Figure5H,J**), indicating that clustering alone does not drive stomatal size increase. To investigate effects of EPF/EPFLs on stomatal size while minimizing patterning defects, we generated FAMAp-driven EPF/EPFL misexpression lines. Late lineage expression of EPFL2 or EPFL6 resulted in a decrease of stomatal complexes (**Figure5I,J**), indicating that ER signaling impacts stomatal size, though it remains unclear how size is normally regulated between leaf surface and tip.

## Discussion

Cell identities are specified during plant embryogenesis but these cells do not become functional until germination. We here show that embryonic maturation of stomata at the cotyledon tip is controlled by ER signaling.

### Defining complex cell states/ levels of maturity based on gene expression

In this study we identify several complex cell states where cells gain aspects of the stomatal maturation program but either arrest or skip other key aspects, resulting in what we describe as partially mature GCs. This represents a re-occurring challenge and opportunity in biology when mutants, treatments or misexpression can result in the creation of new cryptic or mixed cell states. Using single cell transcriptomics these can now more easily be compared to wild-type cell states, such as can be done here with the stomatal lineage^49,55,67^. And while mixed cell identities remain challenging to classify, they do allow us to start untangling the programs that make up cell states^55,68–71^. At the same time, they require updating and refining our definitions of cell states. For GC maturation, a curated set of 586 genes was published^31^ that we here further narrowed down to 214 genes to capture genes as specific as possible to GC maturation with only minimal expression in earlier stomatal stages or other cell types (**TableS3**). For EPF2p::FAMA embryos, our transcriptomic analysis revealed that their stomatal cells express early GC genes but none associated with later stages, likely reflecting a relatively simple arrest during young GC-like cell state (**Figure1G**). In contrast, the tip-located GCs in *er erl1 erl2* embryos express both late GMC genes and late GC genes while lacking key young GC genes (**Figure3F-G**), representing a different cell state.

### Leaf polarity and SCRM activity have limited impact on FAMA-driven GC maturation

To determine what limits FAMA’s ability to drive embryonic maturation, we started from known variables that affect stomatal development. Leaf polarity affects stomatal patterning, with more stomata forming on the abaxial surface^9,72,73^, and developmental speed, with identity acquisition as measured by early stomatal gene expression starting on the embryonic adaxial surface^5^ and postembryonic maturation progressing earlier on the adaxial surface^9^. In EPF2p::FAMA embryos adaxial cells show additional GC features compared to abaxial ones indicating that leaf polarity contributes to stomatal cell state regulation (**Figure1E**). However, this progression is still limited, as transcriptomic analysis revealed that only GC genes associated with young GC stages are upregulated (**Figure1G**).

Another known factor impacting FAMA is the availability of its heterodimerization partners. SCRM-D EPF2p::FAMA embryos, where SCRM activity is abundant, created more stomatal cells including GC-like cells (**Figure1H-I’,S1B-D**). However, no further progression to maturation could be seen, indicating that SCRM presence does not limit embryonic GC maturation potential.

### Absence of ER signaling triggers a unique, partially mature GC state

ER signaling affects stomatal patterning and maturation with loss of ER and its homologs causing an increase in stomatal numbers and increased maturation starting 2 days after germination^50^. We find that germinating *er erl1 erl2* seedlings contain mature-looking GCs 12 hours after being placed in light (**Figure2A,D**) and that these maturing GCs are already present during embryogenesis, starting 6 days after fertilization (**Figure2J**). Any level of ER signaling can prevent embryonic GC maturation as embryos with one functional ER or ERL1 copy lack GCs.

The GCs in *er erl1 erl2* embryos more closely resemble functional complexes compared to those in EPF2p::FAMA embryos: they form pairs of cells with increased PI staining and cell wall thickening at the pore site (**Figure2H,3H**). Comparison between embryonic GC-like cells and ‘regular’ GCs reveals similarities and differences. *er erl1 erl2* embryonic GCs have progressed to and retain features of GMC state as indicated by SDD1 and SLL3 expression (**Figure3A-B**), while transcriptomic analysis reveals expression of GC genes associated with late GC stages (**Figure3F**). While the expression of GC-specific genes indicates acquisition of a GC-like cell state, the absence of many GC genes, including key markers such as SLAC1, GORK, and KAT1/2^27–30^, shows that major aspects of GC state are not present. However, as stomata are a rare cell type, their impact on the overall transcriptome is small and some increases in gene expression can be relevant but not statistically significant such as for FAMA (**Figure3G**).

A key physical GC feature present in *er erl1 erl2* embryos is the start of construction of a central pore. This pore is incomplete but the stage at which it is captured reveals that cell wall reinforcement progresses from the upper and lower surfaces inwards (**Figure3I-K**). How this relates to pore formation during regular GC maturation is not certain but this could be an aspect of pore formation that was easily overlooked in previous TEM imaging, where limited three-dimensional information and off-center sectioning can obscure such features. In addition, our TEM images reveal an absence of vacuoles and presence of plasmodesmata (**Figure3K’’**), features that differ from mature stomatal complexes and further underline lack of cell maturity.

Considering other components of the ER signaling cascade, we find that *yda-11* mutant embryos occasionally have pairs of GC-like cells at the cotyledon tip, confirming that GC maturation is regulated by the canonical ER signaling pathway (**Figure4C-D**). The lower frequency is likely the result of remaining signaling activity. This is hard to eliminate since higher order mutants of ER signaling components are not available or do not produce embryos^34,38,41,60^.

Lack of GC complexes in a high percentage of *er erl1 erl2 fama+/-* embryos indicate that FAMA is required for the maturation processes that occur in the absence of ER signaling. However, only low levels of FAMA appear to be necessary as FAMA transcript levels remain low in *er erl1 erl2* embryos. Low FAMA levels and the differences in embryonic GC states between *er erl1 erl2* and EPF2p::FAMA embryos, led it to hypothesize where combining the two might allow for more extensive maturation. Instead, it leads to both phenotypes co-existing without either GC-type progressing further (**Figure4G-M**). Exactly which ER signaling targets are involved in this remains unknown and they could act separately from stomatal bHLHs since even though SCRM1/2 connect the MAPK cascade to the core stomatal bHLHs^39^, the hyperactive SCRM-D did not mimic the ER signaling phenotype^5^(**Figure1H**).

### Tip-located Guard Cells might be specific Hydathode GCs whose size is regulated by local ER signaling

GCs maturing in *er erl1 erl2* being located at the cotyledon tip informs us on potential regulatory mechanisms. Leaf tips are sites of auxin accumulation^74^ and regional hydathode identity^75^. GC maturation is generally associated with low instead of high auxin signaling^76,77^ and indeed, increased auxin did not increase GC numbers or maturation level (**Figure5A-B**). Thus, auxin does not limit the tip-exclusive GC maturation seen.

The role of hydathode regional identity can not yet be tested as the underlying mechanisms are not understood and markers specific to hydathode GCs could not be identified despite available transcriptomic data63,65,66. Hydathode GCs form via the same pathway as regular GCs but then gain additional features: increasing in size and losing the ability to fully close the pore^65^. We hypothesize that there might be only few truly hydathode GC-specific genes and that instead these cells express size and pore-related genes higher or longer, developing canonical GC features to a drastic level. In that case, the GC genes upregulated in *er erl1 erl2* embryos could be a starting point to identifying hydathode GC-specific factors.

Finally, the large GCs found in embryos and leaf tips of *er erl1 erl2* individuals appear to be the result of local ER signaling rather than stomatal clustering (**Figure5F-J**). In the reverse, increasing EPFL2 or EPFL6 levels in late stomatal lineage cells resulted in reduced stomatal complex size, indicating that EPFL perception by ER affects stomatal size. While the targets and mechanisms remain unclear, ER signaling affects GC size and provides a mechanism for creating leaf-wide differences with ERL1 and ERL2 expression peaking at the embryonic leaf tip(Figure5**D-E**).

Altogether we find that ER signaling uniquely regulates maturation and size of GCs at the cotyledon tip, starting during embryogenesis. However, its regulation does not encompass all facets of the GC maturation, resulting in the formation of partially mature stomatal complexes as during embryogenesis. Our findings now provide a starting point to untangling GC maturation modules and their modifications in embryos and hydathode regions.

## Material and methods

### Plant material and growth conditions

All *Arabidopsis thaliana* lines used in this work are in the Col-0 background. For *er erl1 erl2* plants, we used two independent triple mutant combination segregating for either ER or ERL1. er+/- erl1 erl2 contained er-105 (CS89504 / GABI_182D08), erl1-2^59^, erl2-1 (CS6588 / Wiscseq_DsLoxHs009_09A.2) as previously described in ^59^; er erl1+/- contained er (SALK_066455), erl1 (GK_109G04), and erl2 (GK_486E03) as previously used in ^78^. er -/- erl1 erl2 or er erl1-/- erl2 embryos and seedlings were identified via genotyping or using their characteristic short cotyledon morphology. For er+/- erl1 erl2+/- fama embryos ER signaling homozygotes were likewise identified using their characteristic leaf morphology and small cell clusters.

Newly generated lines and sources of previously reported transgenic lines are listed in the key resources table. Arabidopsis seeds were surface-sterilized, plated on half-strength Murashige and Skoog medium without sucrose and with 0.8% agar. After 2 days of stratification at 4 °C, seedlings were grown for 2-14 days under standard long-day conditions in a Percival growth chamber, model SE-41 (16 hr light/8 hr dark at 150 μmol m2/s and 22°C). Plants were then transferred to soil and grown at standard long-day growth conditions.

### Molecular cloning and plant transformation

An overview of all vectors created for this work is included in the key resources table. Primers used for cloning and genotyping are listed in the Key Resource Table. Constructs for plant transformation were cloned using GreenGate backbones pGGZ003 and pGGZ004^79^. Transgenic plants were generated by floral dip and transgenic seedlings were selected on ½ MS without sucrose with the appropriate antibiotic (15 mg/L phosphinothricin, 7,5 mg sulfadiazine or using FastRed^80^ seed selection, depending on the construct)

### Sample preparation, microscopy and image processing

Mature embryos were dissected from imbibed seeds and stained with modified Pseudo-Schiff Propidium Iodide (mPS-PI) and mounted with Hoyer’s (containing chloral hydrate). Stacks were processed using MorphographX^52^, creating a mesh to mark the tissue boundary onto which to project epidermal signals (∼0-6 μm from the mesh) before segmenting cell areas. Embryos with reporters were stained with Renaissance 2200 after ClearSee destaining^81,82^ and were imaged and processed using FIJI and MorphographX^52^.

### Confocal microscopy

All images were collected via a Leica TCS SP8 system. For mP-PI images, the laser of 561 nm was used for excitation, and the PMT detector collected the emitted light at 570-620 nm. For ClearSee and SR2200 stained images^81,83^, the cell wall was detected via the laser at 405 nm and wavelengths collected at 420-450 nm. Other fluorophores were excited at respective wavelengths, and collected through PMT or Hybrid detectors (HyD) using a 40x water objective or 20x dry objective. Channel bleedthrough was avoided using sequential scans. The excitation (Ex) and emission (Em) wavelengths used for each fluorophores are as follows: mTurquoise-Ex: 458nm, Em: 470-520nm, GFP-Ex: 488nm, Em: 495-515 nm, mCitrine, Venus, and YPet - Ex: 514nm, Em: 515-535nm.

### Sample Preparation for Transmission Electron Microscopy

#### Without Microwaving (Figure 3I-L)

Embryos at the torpedo or bent cotyledon stage were dissected from ovules and immediately transferred into fixation solution containing 2% formaldehyde and 2,5% glutaraldehyde. Samples were post-fixed for 2 h in 1% osmium tetroxide (in water) on ice and 1 h in 1% uranyl acetate at room temperature in the dark. Between each incubation step, samples were washed several times with water. After fixation, cotyledons were dehydrated in a graded ethanol series (75%, 90%) followed by 2x acetone (100%), each for 1h. Dehydrated samples were infiltrated with increasing epoxy resin concentrations (10% for 3 h; 25% for 3 h; 50% for 7 h; 75% overnight; 100% for 7 h and 100% overnight). After resin infiltration, samples were embedded in flat embedding molds and polymerized at 60°C for 2 days. Ultrathin sections (50 nm) were performed at the Leica UC7 ultramicrotome. Sections were contrasted with uranyl acetate (in 50% ethanol) and lead citrate. Analysis of stomata ultrastructure was performed at the Jeol JEM-1400Plus transmission electron microscope operated at 120kV and equipped with a 4K CMOS camera TemCam-F416 (Tietz).

#### With Microwaving

Embryos at the torpedo or bent cotyledon stage were dissected from ovules and immediately transferred into fixation solution containing 2% formaldehyde. After collection, the fixation medium was replaced with fresh fixative consisting of 2% formaldehyde and 2.5% glutaraldehyde.All subsequent preparation steps were carried out using a PELCO BioWave Pro microwave processing system.

Fixation and Post fixation: Primary fixation and post fixation were performed sequentially using 2% formaldehyde and 2.5% glutaraldehyde, followed by 1% osmium tetroxide in H₂O and 1% uranyl acetate in H₂O. For each fixation step, the following microwave protocol was applied: four cycles of 1 min microwave irradiation at 150 W, each followed by a 4 min incubation period without microwave exposure to allow sample cooling. Vacuum was maintained throughout all fixation and post fixation steps.

Samples were washed three times in distilled water for 7.5 min each without microwave treatment, followed by two additional washes of 40 s each at 250 W.

Dehydration and Resin Infiltration: Dehydration was performed using a graded ethanol series (75%, 95%, and 100%), followed by two washes in 100% acetone, each for 40 s at 250 W. Epoxy resin infiltration was carried out using increasing resin concentrations (10%, 25%, 50%, 75%, and two subsequent steps in 100% resin). Each infiltration step consisted of three microwave cycles of 1 min at 250 W, with cooling intervals of 1 min between cycles. The samples were additionally incubated overnight at room temperature in 75% and in the second 100% epoxy resin step.

Polymerization was performed at 60 °C for 48 h.

Sectioning and Imaging: Ultrathin sections (50 nm) were cut using a Leica EM UC7 ultramicrotome, post stained with uranyl acetate and lead citrate, and examined with the Jeol JEM-1400Plus transmission electron microscope operated at 120kV and equipped with a 4K CMOS camera TemCam-F416 (Tietz).

### Seed culture

Ovules were dissected out from the siliques with torpedo stage embryos and were surface sterilized with 70% Ethanol. These were further placed on a seed culture media^84^, including a Vitamins and Plant Preservative Mixture. For auxin treatment, 50 µM auxin dissolved in DMSO was added to the media, and media with just DMSO was used as a control. The ovules were then cultured in a growth chamber under normal growth conditions.

### mRNA sequencing

For the mRNA sequencing experiment, each biological replicate contained a pool of 10 embryos dissected out of the seed coats at two different developmental stages (torpedo or bent cotyledon). Embryos were manually staged based on their morphology. For *er erl1 erl2* mutants, only embryos with small cotyledons characteristic of homozygotes were collected. All samples were harvested at ZT4 (4 h after lights-on). Total RNA was isolated with the RNeasy Micro Kit for Low Biomass RNA Extraction (Qiagen, 73934) and sequencing libraries were prepared with the NEBNext® Single Cell/Low Input RNA Library Prep Kit for Illumina® (New England Biolabs, E6420S) following the manufacturer’s guidelines. Libraries were sequenced in an Illumina NovaSeq X Plus platform with paired-end configuration (150 bp).

### mRNA-seq analysis

Raw reads in FASTQ format were subject to quality control using FastQC^85^. Next, reads were pseudo-aligned to a transcriptomic index of the *Arabidopsis thaliana* TAIR10 genome using kallisto version 0.46.1^86^ with the number of bootstraps set to 30. Quantified gene reads were further analyzed in R version 4.4.2 using the package DESeq2^87^ for normalization, data exploration and differential expression analysis. Before normalization, minimal filtering was applied by removing genes with raw counts of zero in all samples. Next, VSD transformation was used for exploratory data analysis and DESeq2-normalized counts were used to visualize transcript levels of individual genes with ggplot2^88^. Differential expression analysis was performed using DESeq2’s built-in pipeline, based on a Wald test to compare genotypes pairwise per developmental stage. For all tests, genes were only considered differentially expressed when false discovery rate-adjusted p-values were less than 0.05. Standard GO term analysis was performed on upregulated genes using the R package clusterProfiler^89^ to identify enriched biological processes. To specifically test for GC maturation, a list of GC-specific marker genes was manually curated by stringently filtering a previous dataset^31^ against early stomatal expression and expression in other lineages (**TableS3**). To specifically test for activation of FAMA targets, a list of 260 FAMA-regulated genes was obtained from previous datasets^25,54^(**TableS4**). In both cases, enrichment of these gene sets was tested using a hypergeometric test against a background of genes expressed in wild-type embryos. To assess gene expression in post-embryonic stomatal cells, a gene module score was calculated for upregulated genes using the built-in AddModuleScore function from the R package Seurat^90^ and subsequently plotted against an annotated UMAP plot from published single cell RNA-seq data^55^.

### Quantification and Statistical analysis

All images were processed using FIJI software or MorphographX software. Information about the biological replicates, total number of samples analyzed and what n means in each experiment along with the respective statistical test is reported in the figure legends or in Table **S10**. We determined statistical significance using a p < 0.05 cutoff.

## Key Resources

### Chemicals

| Reagent/Resource | Source | Identifier |
| --- | --- | --- |
| Propidium iodide<br>Dye for staining intercellular space in Arabidopsis | Thermo Fisher | P3566 |
| SCRI Renaissance Stain 2200<br>Dye for polysaccharides in Arabidopsis | Renaissance Chemicals | SR2200 |
| 2,4-Dichlorophenoxyacetic acid | Duchefa Biochemie | D0911.0100 |
| Plant Preservative Mixture | Plant Cell Technology | 3808.94 |
| Nitsch Medium | Duchefa Biochemie | N0223.0005 |
| MS media for Arabidopsis | Duchefa Biochemie | M0221.0050 |

### Transgenics

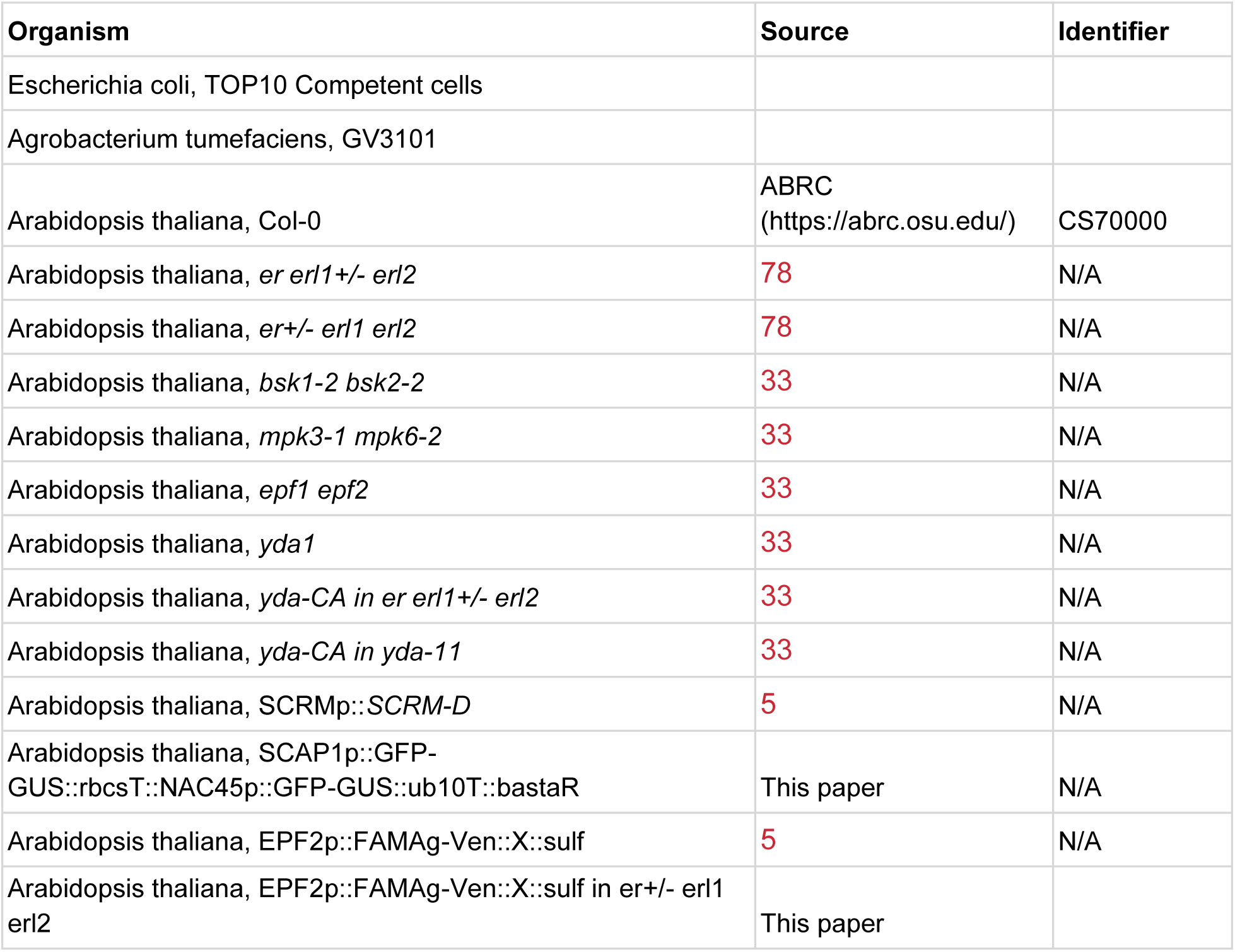
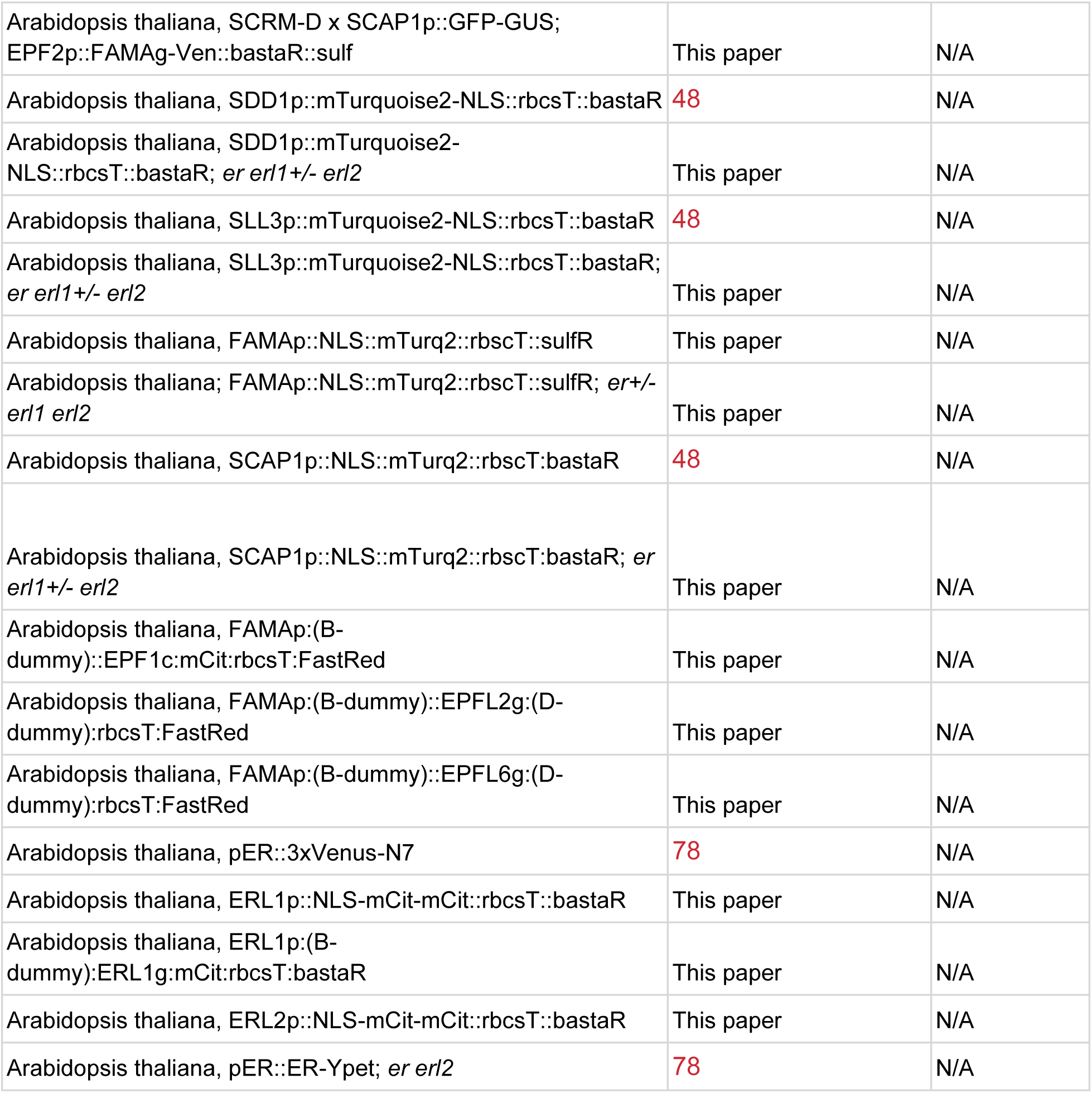

| Organism | Source | Identifier |
| --- | --- | --- |
| Escherichia coli, TOP10 Competent cells |  |  |
| Agrobacterium tumefaciens, GV3101 |  |  |
| Arabidopsis thaliana, Col-0 | ABRC<br>(https://abrc.osu.edu/) | CS70000 |
| Arabidopsis thaliana, *er erl1+/- erl2* | 78 | N/A |
| Arabidopsis thaliana, *er+/- erl1 erl2* | 78 | N/A |
| Arabidopsis thaliana, *bsk1-2 bsk2-2* | 33 | N/A |
| Arabidopsis thaliana, *mpk3-1 mpk6-2* | 33 | N/A |
| Arabidopsis thaliana, *epf1 epf2* | 33 | N/A |
| Arabidopsis thaliana, *yda1* | 33 | N/A |
| Arabidopsis thaliana, *yda-CA in er erl1+/- erl2* | 33 | N/A |
| Arabidopsis thaliana, *yda-CA in yda-11* | 33 | N/A |
| Arabidopsis thaliana, SCRMp::SCRM-D | 5 | N/A |
| Arabidopsis thaliana, SCAP1p::GFP-GUS::rbcS::NAC45p::GFP-GUS::ub10T::bastaR | This paper | N/A |
| Arabidopsis thaliana, EPF2p::FAMAg-Ven::X::sulf | 5 | N/A |
| Arabidopsis thaliana, EPF2p::FAMAg-Ven::X::sulf in *er+/- erl1 erl2* | This paper |  |
| Arabidopsis thaliana, SCRM-D x SCAP1p::GFP-GUS;<br>EPF2p::FAMAg-Ven::bastaR::sulf | This paper | N/A |
| Arabidopsis thaliana, SDD1p::mTurquoise2-NLS::rbcS::bastaR | 48 | N/A |
| Arabidopsis thaliana, SDD1p::mTurquoise2-<br>NLS::rbcS::bastaR; *er erl1+/- erl2* | This paper | N/A |
| Arabidopsis thaliana, SLL3p::mTurquoise2-NLS::rbcS::bastaR | 48 | N/A |
| Arabidopsis thaliana, SLL3p::mTurquoise2-NLS::rbcS::bastaR;<br>*er erl1+/- erl2* | This paper | N/A |
| Arabidopsis thaliana, FAMAp::NLS::mTurq2::rbcS::sulfR | This paper | N/A |
| Arabidopsis thaliana; FAMAp::NLS::mTurq2::rbcS::sulfR; *er+/-*<br>*erl1 erl2* | This paper | N/A |
| Arabidopsis thaliana, SCAP1p::NLS::mTurq2::rbcS::bastaR | 48 | N/A |
| Arabidopsis thaliana, SCAP1p::NLS::mTurq2::rbcS::bastaR; *er*<br>*erl1+/- erl2* | This paper | N/A |
| Arabidopsis thaliana, FAMAp:(B-dummy)::EPF1c:mCit:rbcS:FastRed | This paper | N/A |
| Arabidopsis thaliana, FAMAp:(B-dummy)::EPFL2g:(D-dummy):rbcS:FastRed | This paper | N/A |
| Arabidopsis thaliana, FAMAp:(B-dummy)::EPFL6g:(D-dummy):rbcS:FastRed | This paper | N/A |
| Arabidopsis thaliana, pER::3xVenus-N7 | 78 | N/A |
| Arabidopsis thaliana, ERL1p::NLS-mCit-mCit::rbcS::bastaR | This paper | N/A |
| Arabidopsis thaliana, ERL1p:(B-dummy):ERL1g:mCit:rbcS::bastaR | This paper | N/A |
| Arabidopsis thaliana, ERL2p::NLS-mCit-mCit::rbcS::bastaR | This paper | N/A |
| Arabidopsis thaliana, pER::ER-Ypet; *er erl2* | 78 | N/A |

### Oligonucleotides

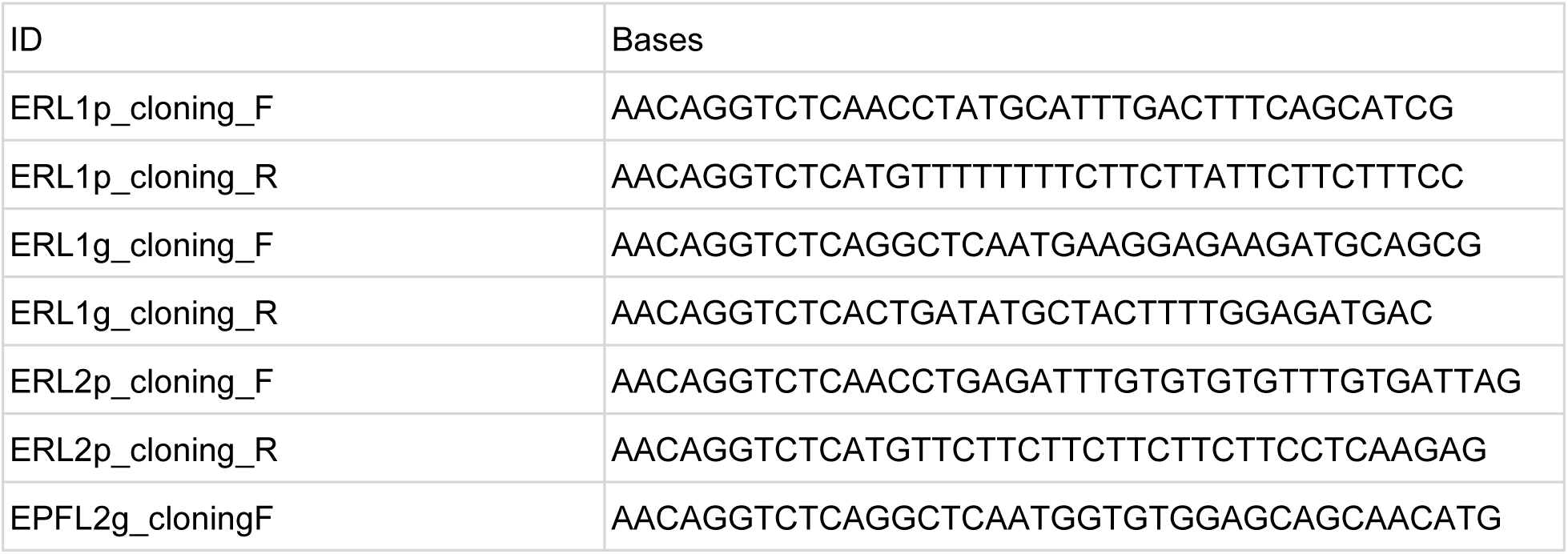
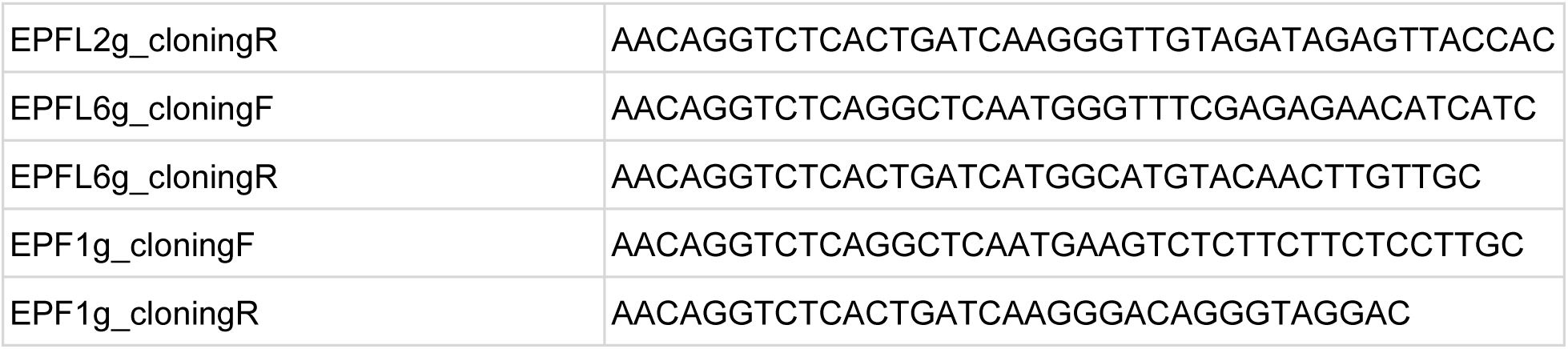

## Acknowledgements

We thank Dominique Bergmann and her lab at Stanford/HHMI where MES created initial reagents for this project and HHMI for providing funding for materials and salary. We would like to thank Rebecca Stahl and Lorenz Henneberg at the ZMBP Microscopy Facility for their support in sample preparation and TEM imaging. We also thank the Bayer lab for valuable discussions and suggestions, as well as for kindly sharing seeds. We thank Tom Denyer for valuable input and advice on RNA-sequencing and Thomas Eekhout and Bert De Rybel for facilitating access to published datasets used here. Finally, this work was enabled by DFG grants to ZMBP microscopy facilities: the SP8 confocal laser scanning microscope and Jeol transmission electron microscope were funded under the instrumentation grants INST 37/819-1 FUGG and INST 37/900-1 FUGG.

## Data and resource availability

The raw data for the RNA-seq datasets generated are being made available through Gene Expression Omnibus (GEO).

## Supplemental Figures

**Figure S1:**
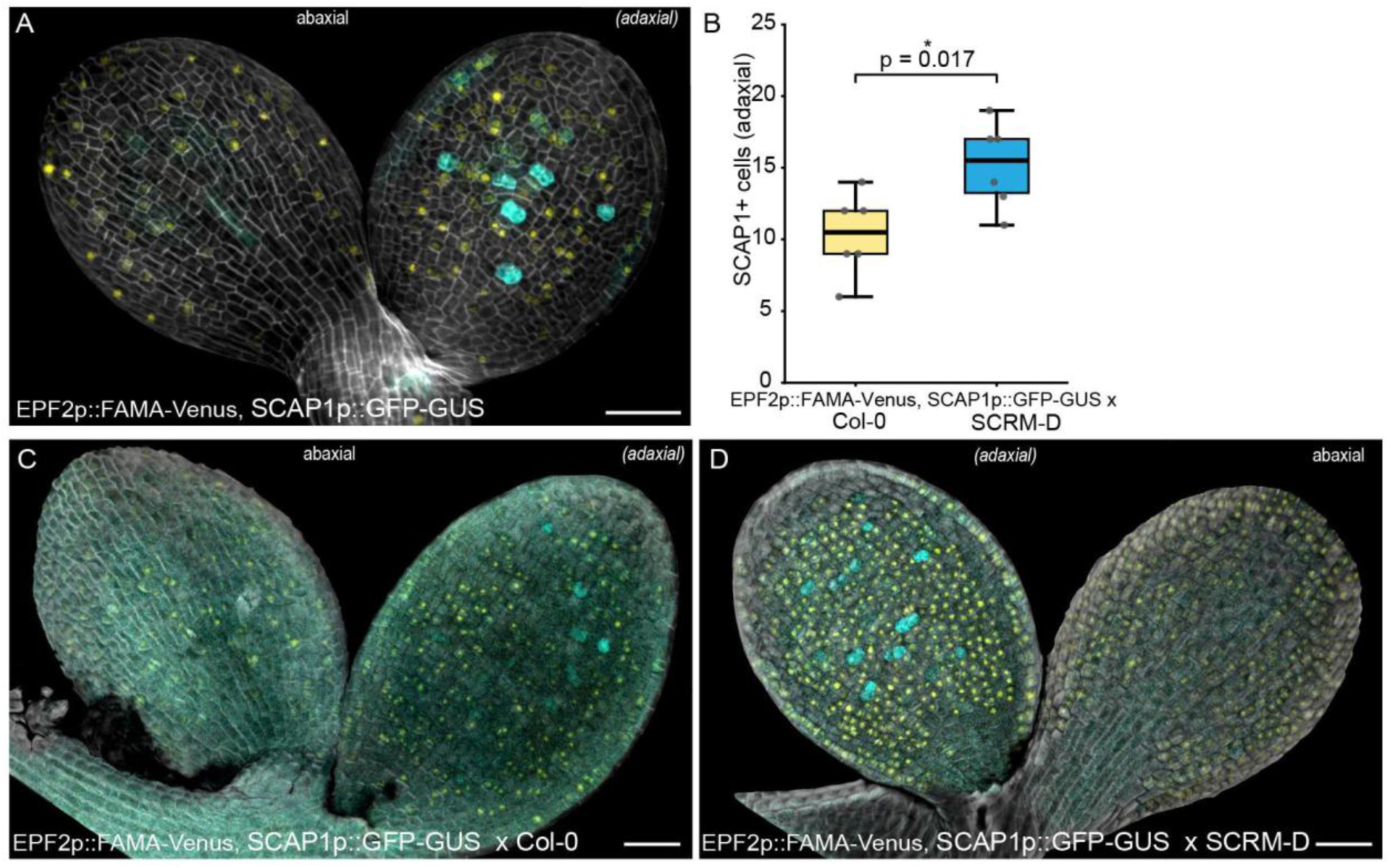
Embryonic FAMA causes partial maturation: (A) SCAP1p::GFP-GUS (in cyan) expression is restricted to the adaxial surface even though EPF2p::FAMA-Venus is present on both epidermal surfaces. (B) EPF2p::FAMA x SCRM-D embryos show a significantly higher number of cells expressing SCAP1 on the adaxial surface than EPF2p::FAMA x Col-0 (n=6 leaves). (C-D) Images showing differences in fluorescence of EPF2p::FAMA-Venus and SCAP1p::GFP-GUS when combined with SCRM-D (D). Scale bars indicate 50 µm.

**Figure S2:**
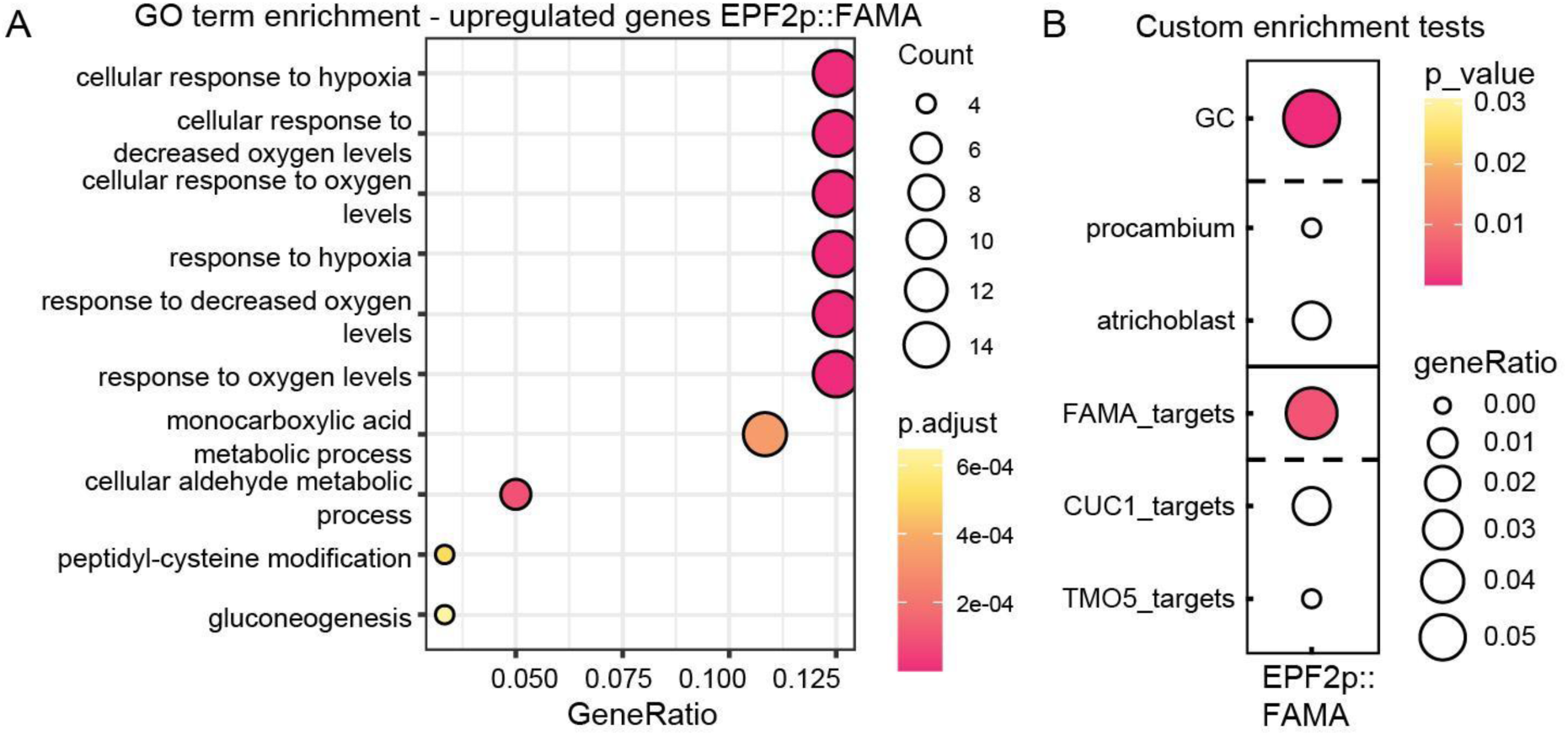
Genes differentially expressed in EPF2p::FAMA embryos are enriched for hypoxia and GC genes. (A) Standard GO term analysis indicates overrepresentation of hypoxia-related genes. (B) Custom enrichment tests show that GC-specific genes and FAMA targets are enriched while other cell type or TF target categories do not show enrichment. (Procambium and Atrichoblast genes from^91^, CUC1 targets^92^, TMO5 targets^93^)

**Figure S3:**
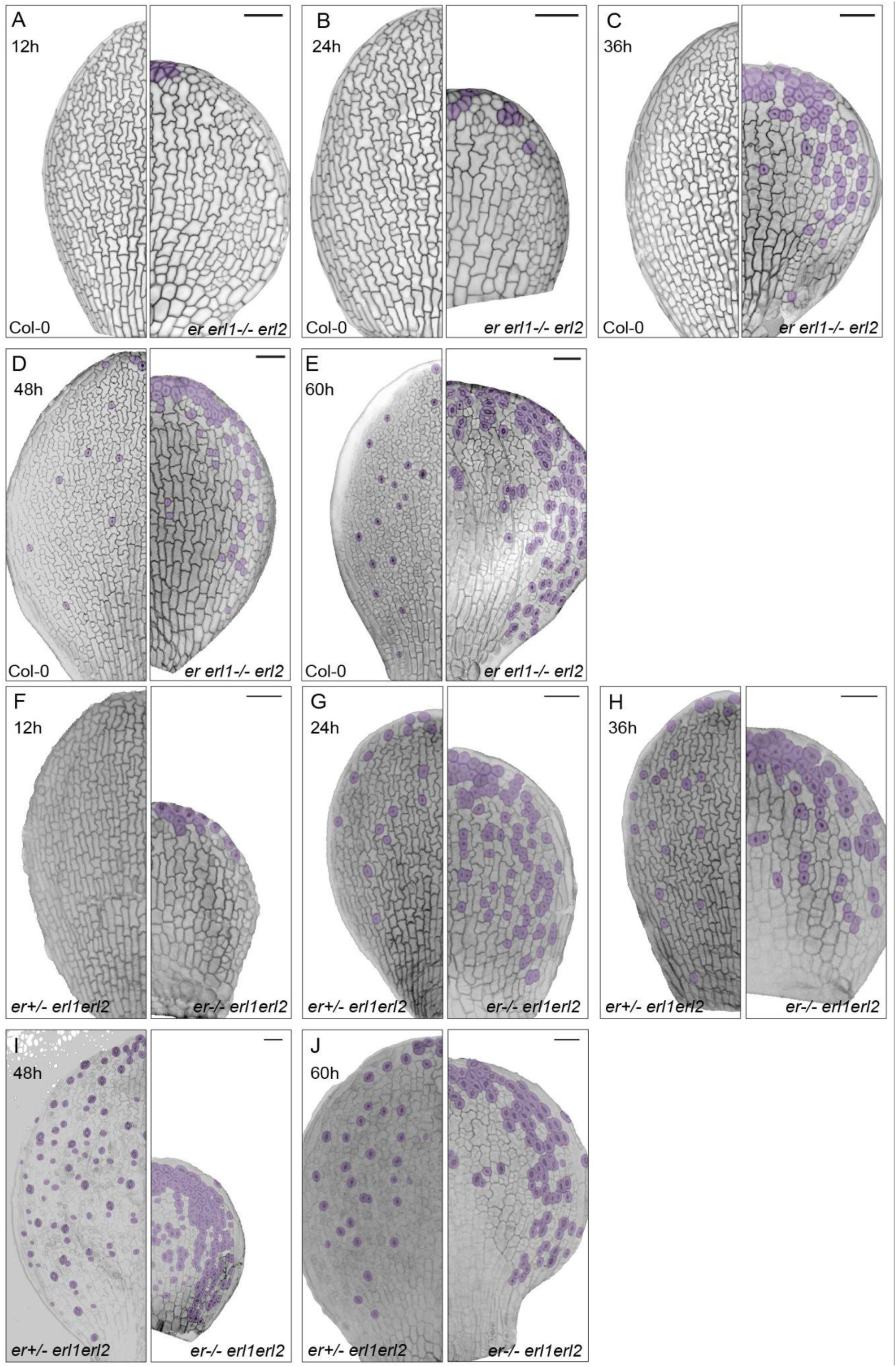
*er erl1 erl2* mutants produce precocious stomata starting during embryogenesis. (A-E) Abaxial epidermal images of Col-0 (left) and *er erl1-/- erl2* (right) germinating seedlings at 12 (A), 24 (B), 36 (C) 48 (D), and 60 (E) hours after being placed into the light. (F-J) Abaxial epidermal images of *er+/- erl1 erl2* (left) and *er-/- erl1 erl2* (right) germinating seedlings at 12 (F), 24 (G), 36 (H) 48 (I), and 60 (J) hours after being placed into the light. Scale bars indicate 50 µm. Purple highlights indicate GCs and GC-like cells.

**Figure S4:**
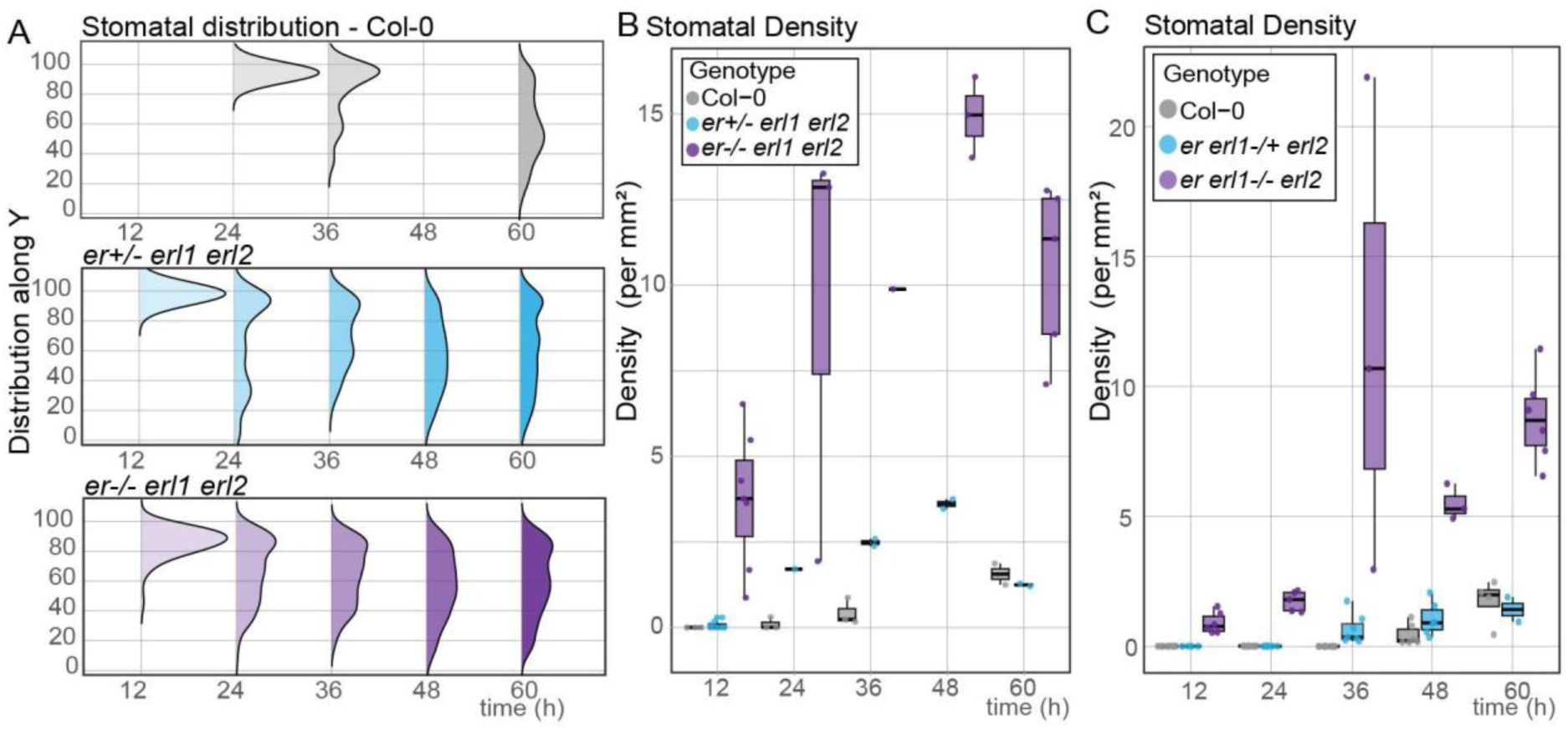
Time series of stomatal density and distribution in WT and *er erl1+/- erl2* cotyledons. (A) Quantification of when and where mature stomata are present upon germination. Y-axis shows the length along the cotyledon, with 0 indicating the base and 100 indicating the tip. Stomatal complexes were counted and the distribution of their Y-coordinates was plotted over time. n=1-8 leaves. (B-C) Quantification of stomatal density of Col-0 and both genotypes of *er erl1+/- erl2* (B) and *er+/- erl1 erl2* (C) upon moving to the light.

**Figure S5:**
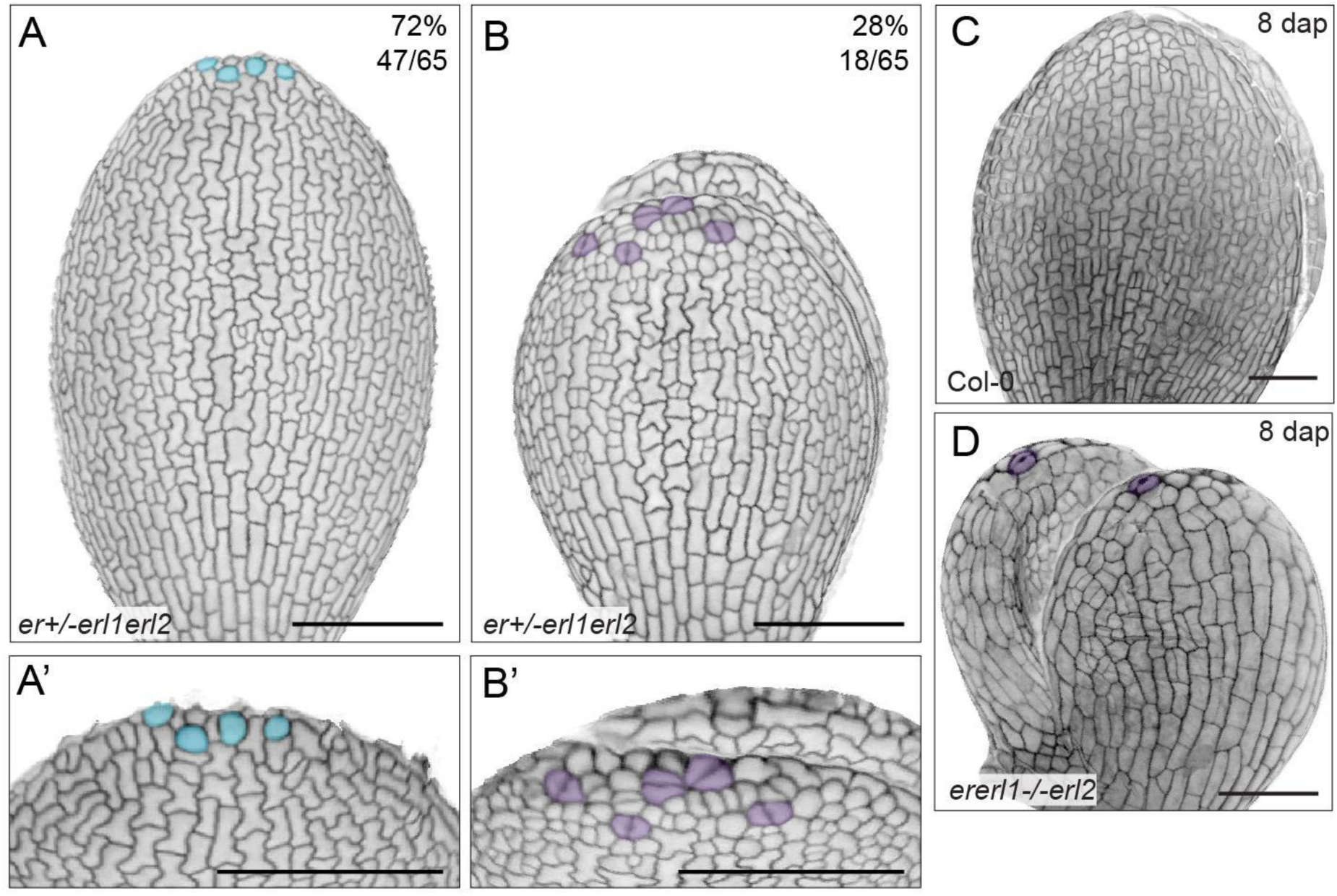
ER mutants show precocious stomatal maturation during embryogenesis. (A) Image of *er+/- erl1 erl2* epidermis, lineage precursors at the tip highlighted in blue and tiled zoom-ins in (A’). (B) Image of *er erl1 erl2* epidermis, maturing GCs at the tip highlighted in purple and tilted zoom-ins in (B’). (C) Image of WT abaxial epidermis, 8 DAP. (D) Image of *er erl1 erl2* abaxial epidermis, 8 DAP, purple cells highlighting maturing GCs, indicating their presence during embryogenesis. Scale bars A-B’ indicate 100 µm, in C-D 50 µm.

**Figure S6:**
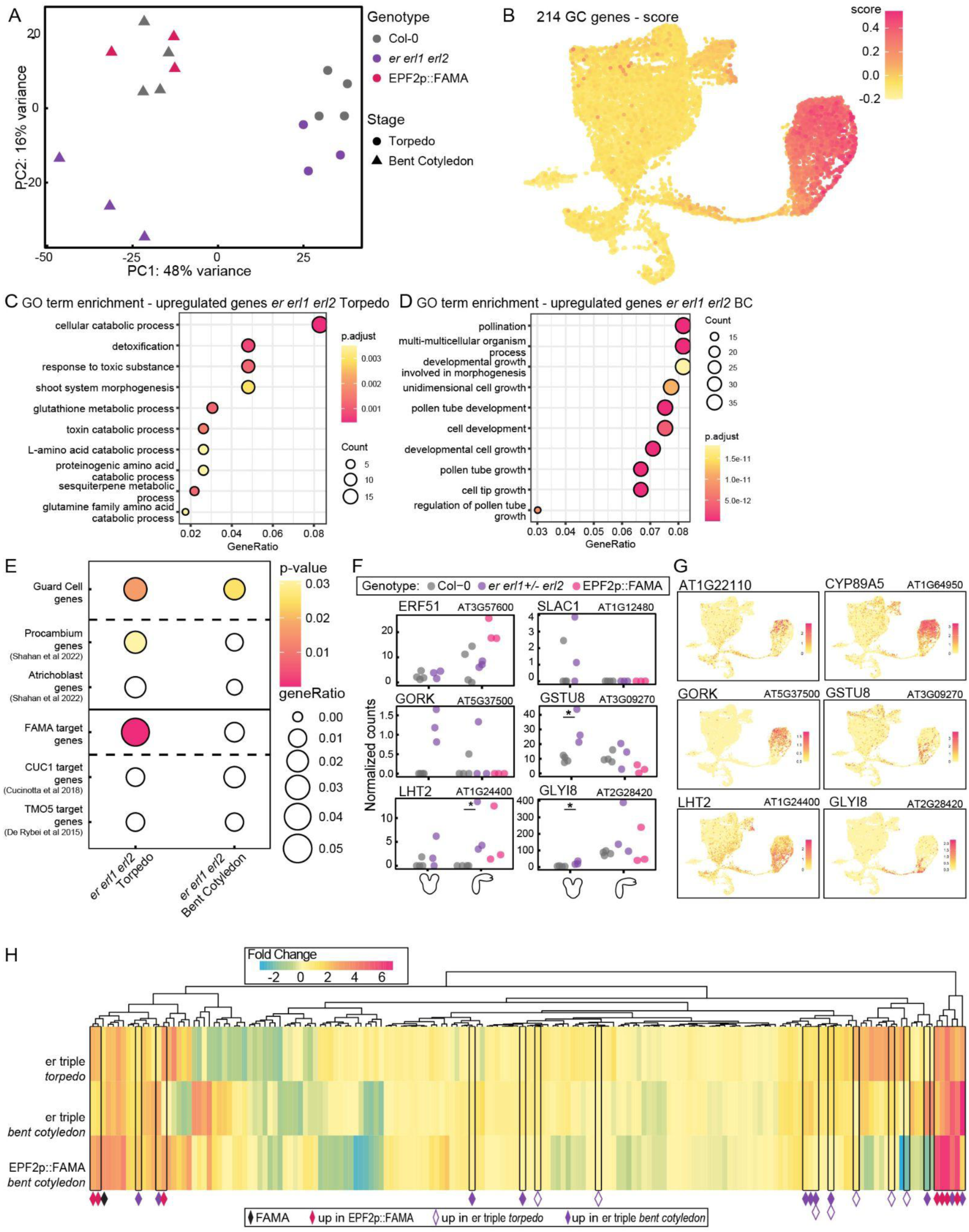
Embryonic transcriptomes reveal the extent of GC maturation in *er erl1 erl2* and EPF2p::FAMA. (A) PCA plot highlighting separation between *er erl1 erl2* Torpedo and BC stages as compared to the respective WT stages and EPF2p::FAMA. (N=3 samples, n=10 embryos per sample) (B) Projection of 214 GC marker genes onto the available scRNA-seq data of the stomatal lineage^55^. (C-D) GO-term enrichment analysis of genes upregulated in *er erl1 erl2* embryos at torpedo stage (C) and bent cotyledon stage (D). (E) Custom enrichment tests for genes upregulated in *er erl1 erl2* embryos including GC-specific genes and FAMA targets. (F) Expression of GC genes in Col-0, *er erl1 erl2,* and EPF2p::FAMA. (G) Stomatal expression of genes from F according to stomatal scRNA-seq data^55^. (H) Heatmap showing which of 214 GC genes are upregulated in *er erl1 erl2* and EPF2p::FAMA.

**Figure S7:**
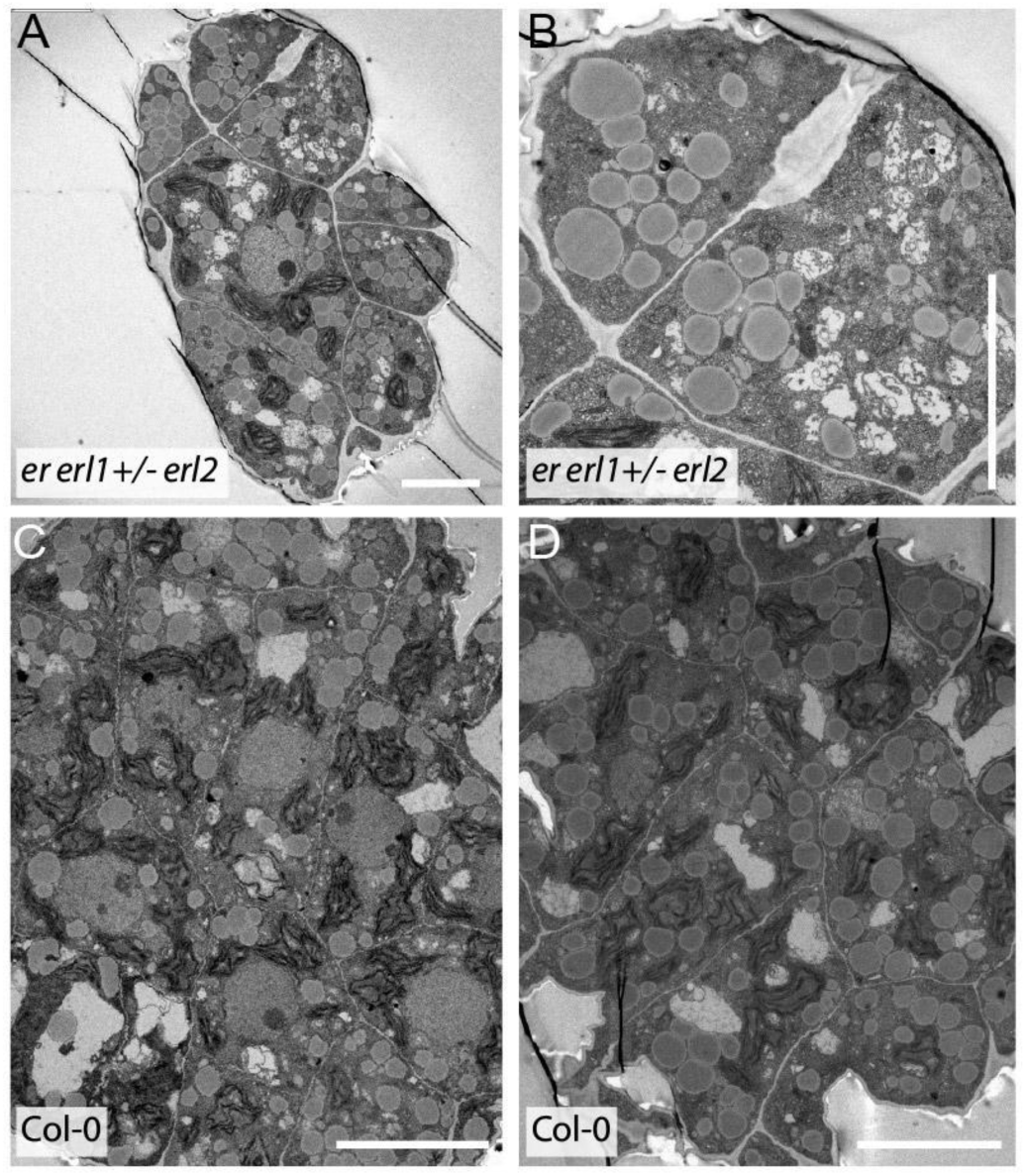
TEM images reveal ultrastructure of embryonic stomata. (A-B) TEM images showing the cross-section of epidermal cells at the leaf tip of a bent cotyledon *er erl1 erl2* embryo with (B) showing detail of cell wall thickening between a pair of GCs. (C-D) TEM images of a WT embryonic cotyledon tips showing no extensive cell wall thickening. Scale bars indicate 5 µm.

**Figure S8:**
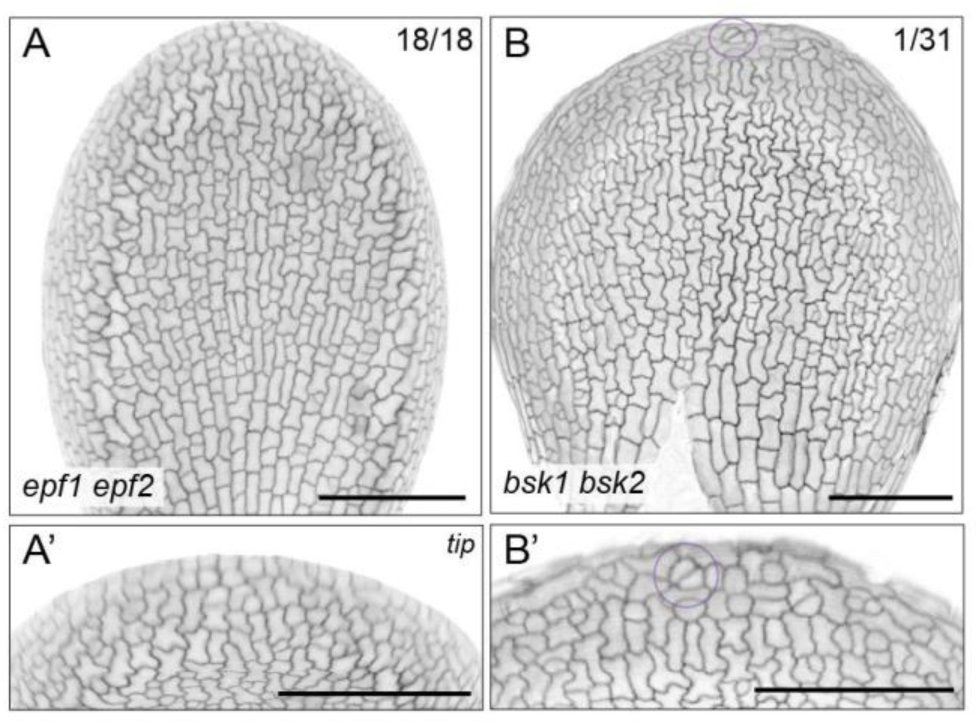
Not all ER signaling mutants show embryonic GCs. (A) Abaxial epidermis of *epf1 epf2* showing enhanced small cell numbers but zoom in image (A’) reveals no maturing GC at the tip (n=18). (B) Only 1 of 31 *bsk1 bsk2* mutant embryos showed a pair of GC-like cells, (B’) highlighting a single GC-like pair at the tip, in the purple circle. Scale bars indicate 100 µm.

**Figure S9:**
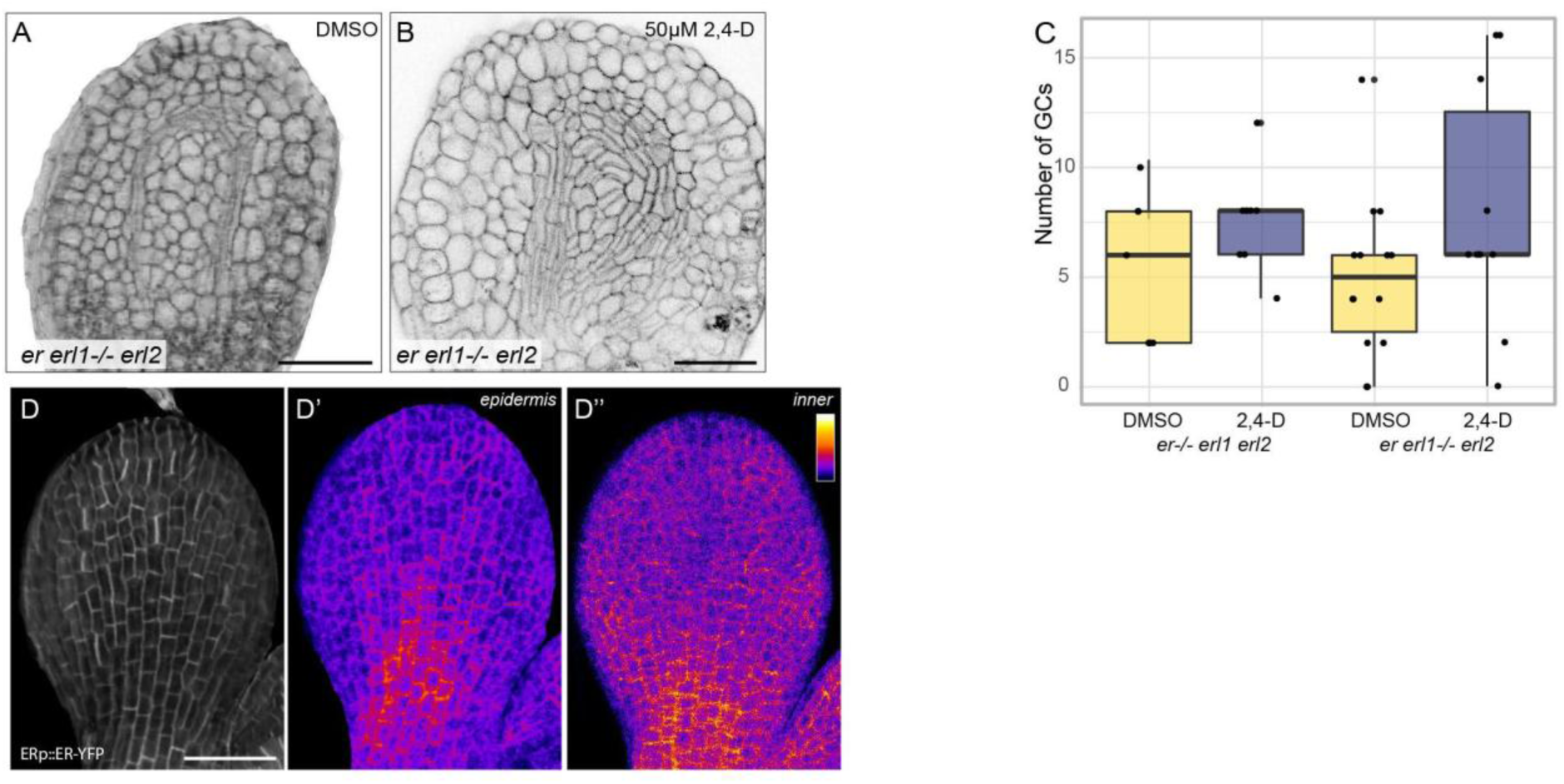
Factors correlating with premature GC maturation at the cotyledon tip. (A-B) Cotyledon vasculature of *er erl1 erl2* embryos treated with DMSO (A) or 50 µM 2,4-D (B) reveals an increase in vascular cell formation, indicating successful auxin treatment. (C) Bar plot showing the number of GCs at the tip of the cotyledon for DMSO vs 2,4-D treated embryos, for both er-/- erl1 erl2 and er erl1-/- erl2 genotypes. (D) Localization of ER during embryogenesis showing that (D’) while it appears homogeneously present throughout the epidermis, (A’’) ER seems to be lower in the hydathode mesophyll cells. Scale bars indicate 50 µm.

